# Field phenomics reveals genetic variation for transpiration response to vapor pressure deficit in sorghum

**DOI:** 10.1101/2023.06.23.546345

**Authors:** Rubí Raymundo, Xu Wang, Terry Felderhoff, Sarah Sexton-Bowser, Jesse Poland, Alexander E. Lipka, Geoffrey P. Morris

## Abstract

Drought adaptation for water-limited environments relies on traits that optimize plant water budgets. Limited transpiration (LT) reduces water demand under high vapor pressure deficit (VPD) (i.e., dry air condition), conserving water for efficient use during the reproductive stage. Although studies in controlled environments report genetic variation for LT, confirming its replicability in field conditions is critical for developing water-resilient crops. Here we test the existence of genetic variation for LT in sorghum in field trials and whether canopy temperature (T_C_) is a surrogate method to discriminate this trait. We phenotyped transpiration response to VPD (TR-VPD) via stomatal conductance (g_s_), canopy temperature (T_C_) from fixed IRT sensors (T_Cirt_), and unoccupied aerial system thermal imagery (T_Cimg_) in 11 genotypes. Replicability among phenomic approaches for three genotypes revealed genetic variability for TR-VPD. Genotypes BTx2752 and SC979 carry the LT trait, while genotype DKS54-00 has the non-LT trait. T_C_ can determine differences in TR-VPD. However, the broad sense heritability (*H^2^*) and correlations suggest that canopy architecture and stand count hampers T_Cirt_ and T_Cimg_ measurement. Unexpectedly, observations of g_s_ and VPD showed non-linear patterns for genotypes with LT and non-LT traits. Our findings provide further insights into the genetics of plant water dynamics.

## INTRODUCTION

Water scarcity is the main threat to agriculture in semi-arid regions. The increasing risk of crop failure in such areas is exacerbated by recurrent droughts, which vary over space and time (Yuan et al., 2019). Thereby ensuring demand for food and fiber in a world facing climate variability and climate change is essential. Adapting with development of climate-resilient crops is a promising strategy to cope with water scarcity (Lorite et al., 2018). However, the nature of drought adaptation traits is complex (Blum, 2011; Cooper and Messina, 2022) since transpiration is driven by many environmental factors. Hence, dissecting these traits requires understanding plant response to each variable (Figure 1A). Depending on the soil water status, plants sense water stress in leaves or roots, causing partial stomatal closure (Sampaio Filho et al., 2018). Water stress in leaves is triggered by vapor pressure deficit (VPD) (Grantz, 1990), while water stress in root tissue is stimulated by soil water deficit (Turner et al., 1985).

**Figure 1.**
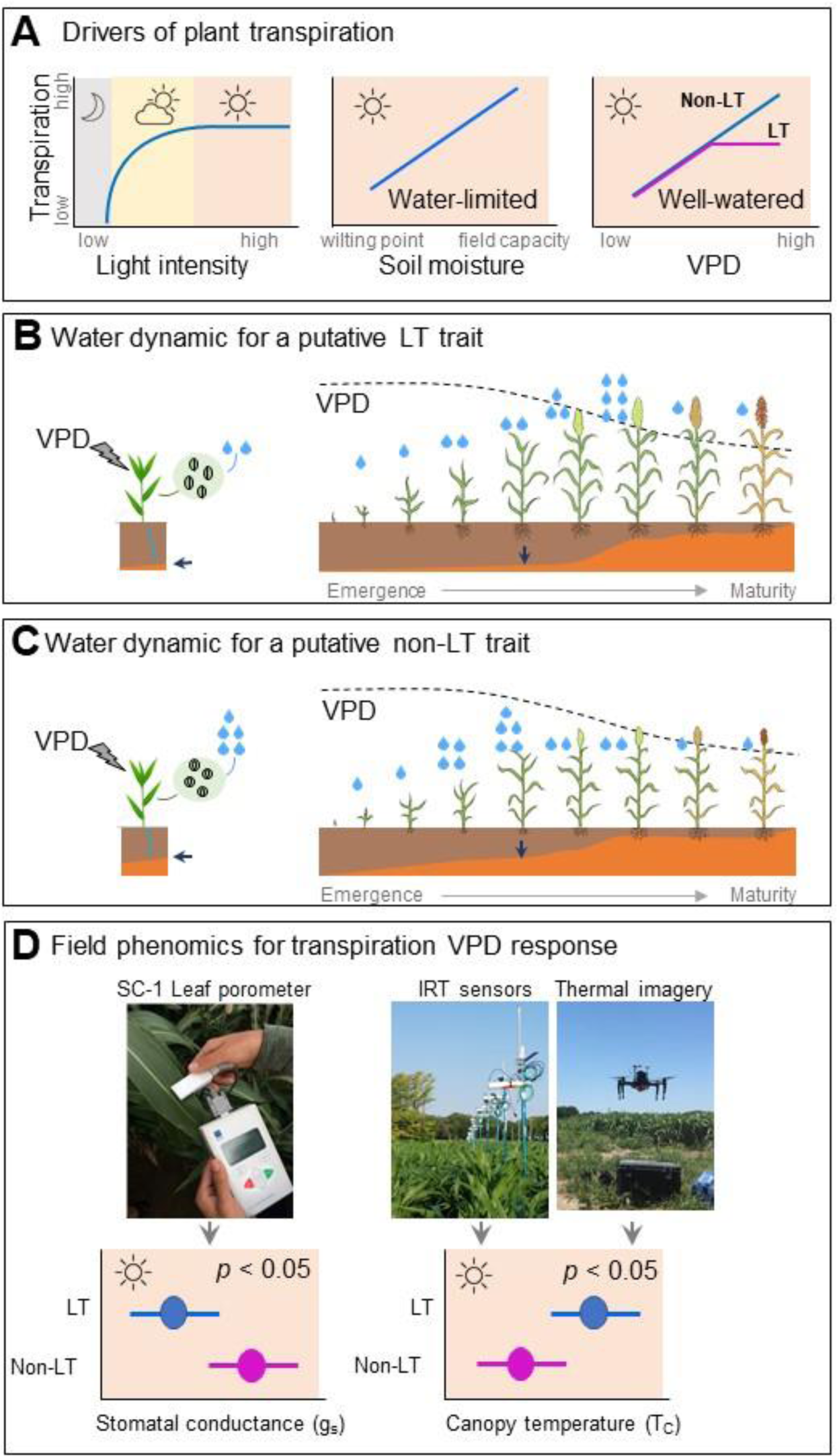
Drivers of plant transpiration, the hypothetical effect for transpiration response to VPD traits during a growing season, and predictions of field phenotyping for transpiration response to VPD traits. A) Transpiration responses to light intensity represented with an asymptotic function (Q), transpiration response to soil moisture represented with an linear function (SW), hypothesis for the transpiration response to VPD for a putative non-LT (linear) and LT trait (breakpoint). Light intensity regulates stomatal response at night ( ), from sunrise to noon ( ), or during any overcast period or day ( ). LT, as a response to light saturation occurs around noon ( ). In this period, transpiration is modulated by SW and VPD in water-limited and well-watered conditions, respectively. B and C) Hypothesis of the effect of the putative LT and non-LT trait in water dynamics during the growing season. Leaves of a genotype with a putative non-LT trait are insensitive to high VPD (low humidity) and keep the stomata open. Leaves of a genotype with putative LT trait perceive a high VPD (low humidity), causing partial stomatal closure. Blue arrows indicate the effect of the LT on soil moisture during the growing season. D) Phenomic approaches to discriminate TR-VPD under well-watered conditions phenotyped via stomatal conductance (g_s_) and canopy temperature (T_Cirt_ and T_Cimg_).

Limiting transpiration (LT) during periods of high VPD would cause partial stomatal closure and redistribute plant use of water within the growing season (Sinclair et al., 2005). Simulation studies revealed that unlike the non-limited transpiration (non-LT) trait, the LT trait reduces water uptake during the vegetative stage, saving soil moisture for grain filling (Figure 1B and 1C). Studies in controlled environments reported genetic variability via breakpoint (LT) and linear (non-LT) transpiration response to VPD (TR-VPD). Under the hypothesis that breakpoint thresholds are heritable, breeding programs could develop commercial hybrids and allocate them to geographies that match the hybrid’s breakpoint TR-VPD to increase water productivity. However, studies were pseudo-replicated, and some studies reported the lack of reproducibility between controlled environments and field trials varied for different crops (Gilbert et al., 2011; Guiguitant et al., 2017; Shekoofa et al., 2014). Thus, replicability of these findings requires rigorous field testing since TR-VPD phenotypic characterization to-date is limited to controlled environments, and most genotypes were not tested in independent field trials (Fletcher et al., 2007; Gilbert et al., 2011; Guiguitant et al., 2017; Schoppach and Sadok, 2013; Shekoofa et al., 2014; Yang et al., 2012).

Phenomic approaches, such as whole plant transpiration and gas exchange, to discriminate TR-VPD are expensive, time-consuming, and limited for few genotypes (Kirkham, 2014). To accelerate selection, breeding programs require large-scale high-throughput plant phenotyping (HTPP) methods, which can effectively replicate the phenotype identified via traditional methods. Canopy temperature (T_C_) is a proxy to estimate plant water stress (Belko et al., 2013; Jackson et al., 1981; Jones et al., 2009) due to its relationship with stomatal response (Deery et al., 2019; Leinonen and Jones, 2004), which is grounded in the theory of energy balance (Jackson et al., 1981). Studies under well-watered conditions suggest that warmer T_C_ and low transpiration can save soil moisture (Pinter et al., 1990). Further, it was hypothesized that genotypes with warmer T_C_ could yield better in drought environments (Pinter et al., 1990). While some studies suggest T_C_ as a reliable approach to dissecting water use traits (Anderegg et al., 2021; Belko et al., 2013; Mutava et al., 2011) other studies indicate that T_C_ can be an artifact of canopy architecture or micro-environmental variation (Prashar and Jones, 2014).

Sorghum is grown and adapted to water limitations. Despite this advantage, its full potential still needs to be explored to identify traits contributing to water productivity (Borrell et al., 2014; Vadez et al., 2014). This study aims to validate and identify the TR-VPD in field trials in the target environment of the sorghum-producing region of the United States, predominantly overlaying the semiarid Great Plains. We hypothesize that sorghum has a genetic variation for TR-VPD under field conditions. Under this hypothesis, our results will be comparable to previous studies. First, we expect low stomatal conductance (g_s_) and high T_C_ for genotypes with the LT trait and the opposite for genotypes with the non-LT trait (Figure 1D). Second, we expect observations to fit breakpoint and linear TR-VPD for genotypes with LT and non-LT traits, respectively. Our second hypothesis is that T_C_ is a surrogate method to discriminate TR-VPD. Under this hypothesis, first we expect a significant negative correlation between g_s_ and T_C_. Second, no significant correlation between T_C_ and canopy architecture traits.

## MATERIALS AND METHODS

### Design and management of field experiments

This study tested the TR-VPD of various genotypes in Ashland Bottoms (AB), Kansas and Greeley (GR), Colorado (Figure 2A). Based on the PRISM dataset (https://prism.oregonstate.edu/), the maximum VPD in summer in AB is 2.5 and 3.5 in GR respectively, corresponding to subhumid and semiarid climates (Figure 2B and 2C). Field trials were planted in the summer seasons of 2019, 2020, and 2021 in AB and GR. The experiment included genotypes with putative TR-VPD and commercial hybrids (Table 1). In 2019 and 2020, the field trials comprised 10 genotypes and only 6 in 2021 (Table 1). Fewer genotypes were planted in 2021 because the expected TR-VPD (Figure 1D) in AB19 and AB20 matched only for genotypes DKS54-00, Tx430, BTx2752 and SC979.

**Figure 2.**
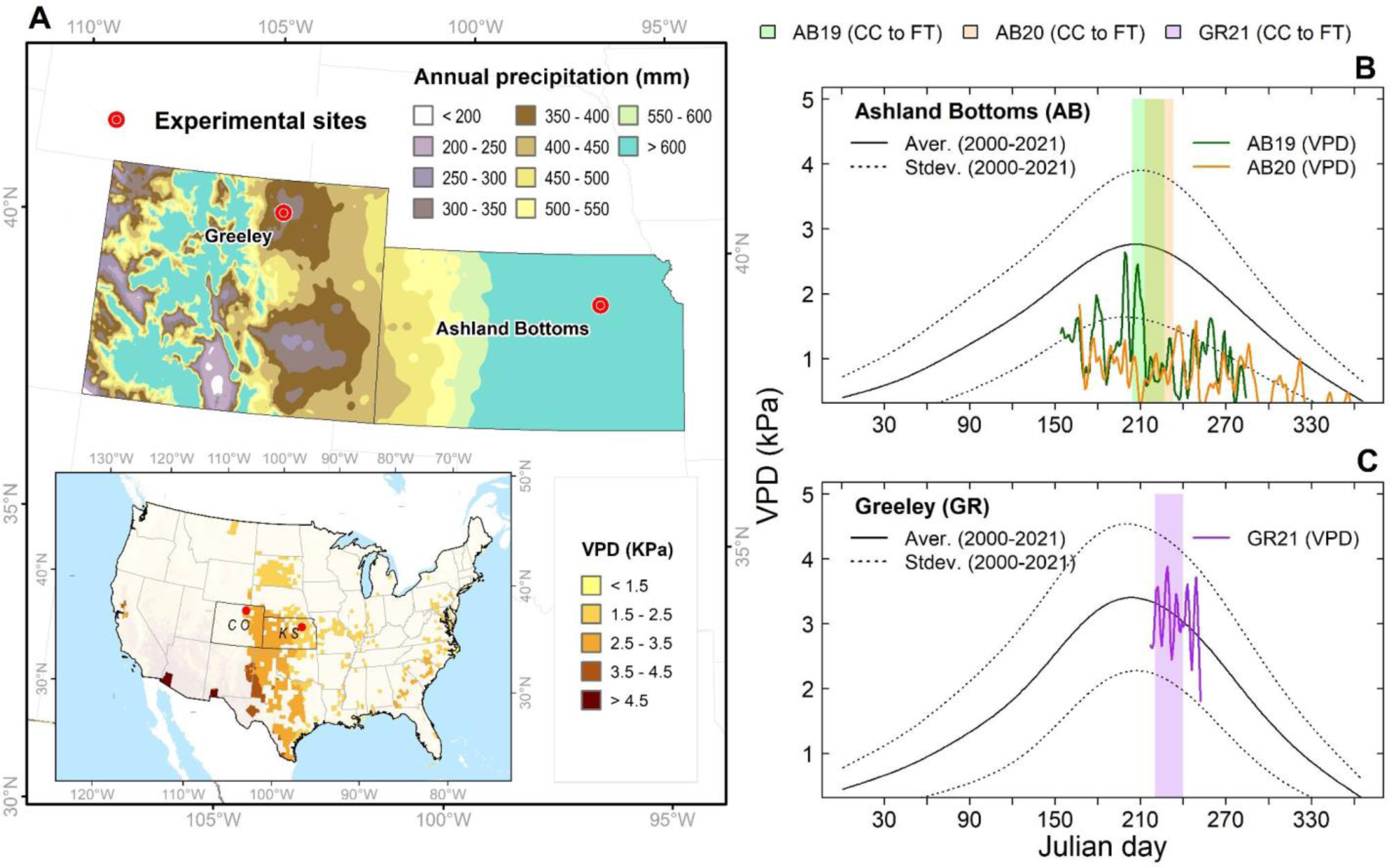
Study system to dissect transpiration response to VPD in sorghum germplasm. A) Spatial variability of annual precipitation and maximum VPD in Kansas and Colorado. Precipitation and VPDe deficit information for sorghum-producing areas were acquired from the PRISM Climate Group (https://prism.oregonstate.edu/). B and C) The annual trajectory of daily maximum VPD from 2000 to 2021 and daily maximum VPD for field experiments in Ashland Bottoms (AB19, AB20) and Greeley (GR21). Maximum daily VPD at each site were acquired from the PRISM Climate Group (https://prism.oregonstate.edu/). Maximum VPD for each experiment was recorded with the ATMOS41 weather station. Information of AB21 not included in Figure 2B.

**Table 1.** Summary of hypotheses on transpiration response to VPD classification for sorghum genotypes reported by previous studies. Stomatal conductance (g_s_) and canopy temperature (T_Cirt_) are proxies of transpiration response to VPD evaluated in field experiments from 2019 to 2021 in Kansas and Colorado (see Table S1). Information on canopy temperature was not collected in AB21.

| Genotype | Maturity | Inferred VPD response | | | $g_s$ | | | $T_{Cirt}$ | | |
| --- | --- | --- | --- | --- | --- | --- | --- | --- | --- | --- |
|  |  | Growth chambers |  | Field | (years) |  |  | (years) |  |  |
|  |  | 31 °C | 37 °C |  | 19 | 20 | 21 | 19 | 20 | 21 |
| SC803 | Med | LT<sup>a</sup> | non-LT<sup>d</sup> | LT<sup>e</sup> | + | + |  | + | + |  |
| SC979<sup>1</sup> | Med | LT<sup>a</sup> | non-LT<sup>d</sup> | - |  | + | + | + | + | + |
| BTx2752<sup>2</sup> | Late | LT<sup>a</sup> | non-LT<sup>d</sup> | LT | + | + | + | + | + | + |
| BTx623 | Late | LT<sup>ab</sup> | non-LT<sup>d</sup> | - |  | + |  | + | + |  |
| BTx642 | Late | LT<sup>a</sup> | non-LT<sup>d</sup> | LT<sup>e</sup> |  | + |  |  | + |  |
| Macia<sup>3</sup> | Late | LT<sup>a</sup> | LT<sup>d</sup> | - |  | + |  | + | + |  |
| Tx7000 | Med | LT<sup>b</sup> | - | - |  | + |  |  | + |  |
| SC1345<sup>1,3</sup> | Early | non-LT<sup>a</sup> | - | - |  |  |  | + |  |  |
| Tx430<sup>3</sup> | Late | non-LT<sup>a</sup> | - | - |  | + | + | + | + | + |
| DKS28-05<sup>4</sup> | Early | non-LT<sup>a</sup> | - | - |  |  |  | + |  |  |
| DKS54-004<sup>4</sup> | Late | non-LT<sup>a</sup> | - | non-LT<sup>e</sup> | + | + | + | + | + | + |
| 84G62<sup>4</sup> | Late | NN | NN | NN | + | + | + | + | + | + |
| ADVG2275 | Med | NN | NN | NN |  |  | + |  |  | + |

NN not known (Commercial checks)
- not available.
+ genotypes evaluated in each experiment.
Germplasm characteristics: <sup>1</sup> Has a Tx430 recombinant inbred line (RIL) family, <sup>2</sup> Elite genotype (G2P
Bridge female tester), <sup>3</sup> Nested association mapping parent (NAM) Parent, and <sup>4</sup> Commercial hybrids.
References: <sup>a</sup> (Gholipour et al., 2010), <sup>b</sup> (Choudhary et al., 2013a), <sup>d</sup> (Riar et al., 2015), and <sup>e</sup> (Shekoofa et al., 2014).

Experiments were planted under a randomized complete block with four replications in 2019 and 2020. In 2021, experiments comprised two water treatments (irrigated and rainfed) planted in a split-plot design. In these experiments, the rainfed treatment tested genotypes’ performance under drought conditions. Experiments were irrigated to isolate transpiration driven by atmospheric VPD and avoid confounding effects due to low soil moisture (Turner et al., 1985; Zhang et al., 2018). In AB19 and AB20, experiments were irrigated daily from full canopy cover to flowering time using drip lines. In GR21, the experiment was irrigated every week from planting to hard dough stage. Irrigations at this site aimed to fill soil moisture up to 100%.

### Atmospheric and soil variables

In each experiment, a weather station (ATMOS 41, METER Group, Inc., USA) recorded temperature, relative humidity, solar radiation, precipitation, wind speed, and VPD at 15-minute intervals. In 2019, soil moisture sensors (TEROS 10, METER Group, Inc., USA) were installed at two depths (15 cm and 30 cm) in a plot of genotype 84G62. In 2021 three soil moisture sensors were installed on three experimental units at 30 cm. In Colorado (GR21), the seasonal precipitation was 107 mm; in Kansas, it ranged from 338 (AB21) to 520 mm (AB19). Over three years of assessment and across all locations, the maximum VPD ranged from 1.1 (AB) to 4.2 kPa (GR), and soil moisture at 30 cm ranged from 0.20 m^3^ m^-3^ to 0.31 m^3^ m^-3^. Experiments with the lowest and highest VPD were AB19 and GR21, respectively (Figure 2B and 2C). General characteristics of the field site, crop management, and environmental conditions for each season are listed in Table S2.

### Stomatal conductance phenotyping

A steady-state SC-1 leaf porometer measured (METER Group, Inc. USA) the water diffusion in the stomatal cavity (Kirkham, 2014). Abaxial stomatal conductance (g_s_) was taken on the third fully expanded sunlit leaf on three representative plants per plot during the vegetative stage. Information was taken between 12:00 to 16:00 hrs. Evaluations were done without cloud cover to ensure observed transpiration was due to atmospheric VPD and not poor light quality. In GR21, stomatal conductance was evaluated 24 hours after irrigation. Environmental conditions (Table S2) for each evaluation were pulled from the ATMOS 41 weather station.

### Canopy temperature phenotyping

Canopy temperature (T_C_) was recorded via proximal sensing by infrared thermometer (IRT) sensors fixed on the ground and remote sensing by thermal cameras carried by an unoccupied aerial vehicle (UAV). A network of fixed IRT sensors wirelessly transmitted T_C_ measurements to a data logger (Dynamax Inc.). T_C_ collected by IRT sensors (T_Cirt_) was recorded in three replications in AB19 and two replications in AB20 and GR21. Each IRT sensor has a 20-degree field of view and covers a circle area on the target with a diameter-to-sensor distance ratio of 1:3 and a 0.5 °C accuracy over a temperature range of 0 °C to 50 °C. In this study, IRT sensors were installed at around 1.5 m above the crop canopy, hence covering circle areas on the canopy with a 0.5m^2^ diameter. Measurements of T_Cirt_ were analyzed from full canopy cover to pre-flowering time to avoid any confounding effect of soil or pollen/flower temperature. For quality control, records of T_Cirt_ (minutes) were aggregated at an hourly time step, and the time series was plotted for visual inspection as a quality control.

A UAV high-throughput plant phenotyping (HTPP) system, including a quadcopter (Matrice 100, DJI, Shenzhen, China) and thermal camera (VUE Pro R, FLIR, USA), were integrated to collect thermal images for T_C_ extraction in AB19, AB21, and GR21 field trials. Raw thermal images were collected under different heat conditions within the same day (i.e., morning, noon, and afternoon) to monitor T_C_ variation between genotypes (Table S2). Aerial image overlap rate between two geospatially adjacent images was set to 80% both sequentially and laterally to ensure optimal orthomosaic photo stitching quality. All data collection flights were operated at 35m above ground level at 3.5 m s^-1^. After taking off, the quadcopter will hover at a waypoint outside the field trials for at least 2 min before real image collection, allowing the thermal camera to self-calibrate under relatively stable ambient air condition. Multiple ground temperature reference panels (Figure 3B) for thermal imaging calibration were placed beside the field trials at least 20 min. before the real data collection. Ground reference temperature values were measured and recorded in a data logger (CR3000, Campbell Scientific, USA). A semi-automated image processing pipeline was used to generate field orthomosaic photos and trait data extraction (Wang et al., 2020). Plot-level T_Cimg_ is the median of all T_C_ values extracted within each manually-generated plot boundary. The two T_C_ methods were compared using the RMSE (Wallach et al., 2014).

**Figure 3.**
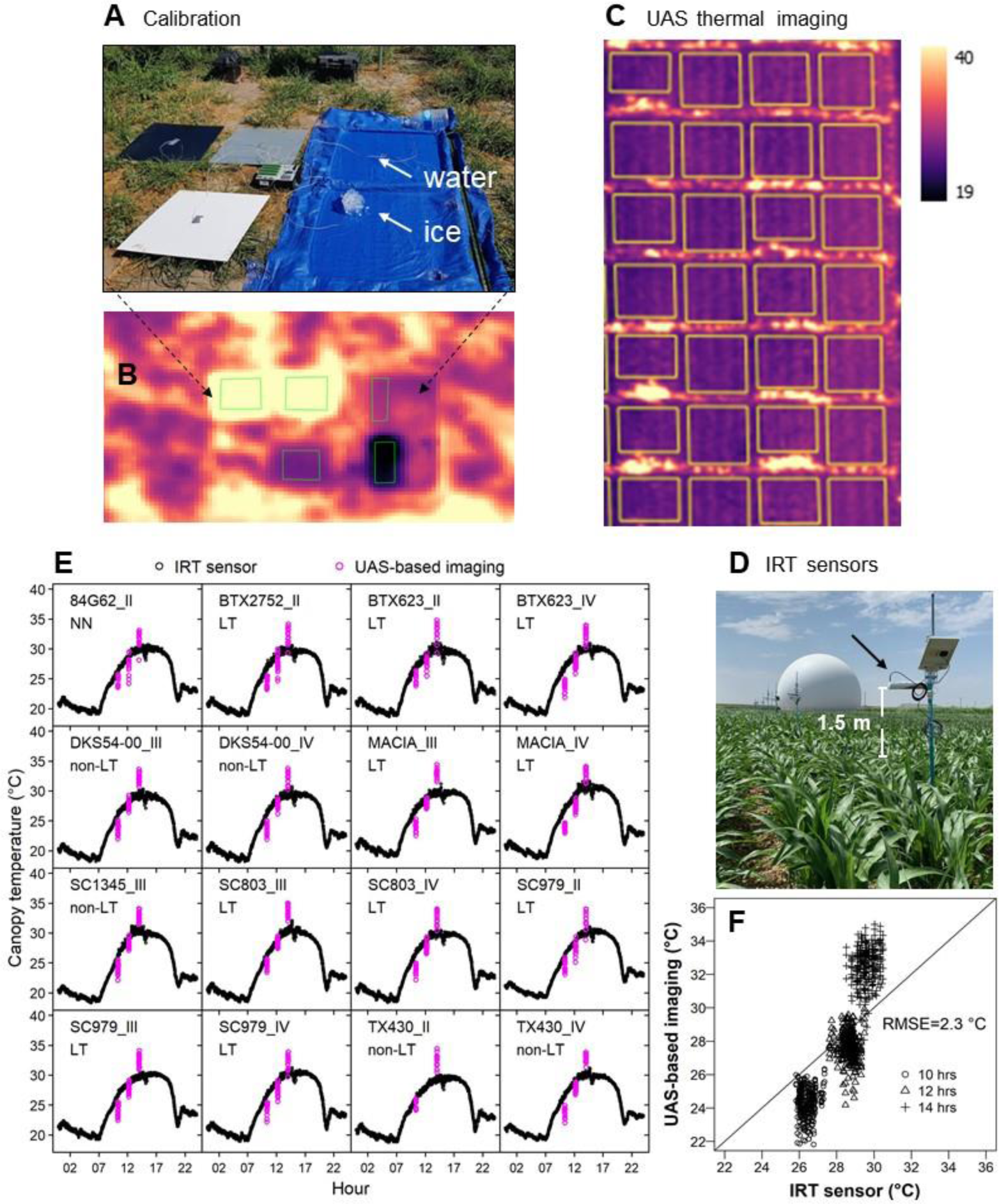
Approach for comparison of sources for canopy temperature (T_C_) in AB19. A) Multiple ground temperature reference panels, including stainless steel boards painted in white, gray, and black, flat square containers filled with water and mixture of ice and water in AB19. B and C) Thermal image reflecting estimated T_C_ in AB19. D) IRT sensors in GR21. Each sensor was located on top of the canopy at a 1.5-meter distance. E) Trajectory of T_C_ obtained with IRT sensors (black symbol) and UAS-based imaging (magenta symbol) in 16 sorghum plots in Ashland Bottoms on July 26, 2019 (AB19). Black symbols represent T_Cirt_ every minute during 24 hours, and magenta symbols represent T_Cimg_ every two seconds during a flight time of 12 minutes. UAS-based imaging was collected in motion at 35 meters above ground level. F) Comparison of T_C_ obtained with IRT sensors and UAS-based imaging in 16 sorghum plots at 10:00, 12:00, and 14:00 hours in Ashland Bottoms on July 26, 2019 (AB19).

### In-season field field phenotyping

After seedling emergence, crop establishment was scored via plant density (Figure S6). In AB19, the plant density ranged from 19 to 25 plants m^-2^. Even though seeding rate emergence was similar in AB20 and GR21 (Table S2), plant density differed in both experiments. For instance, in AB20 and GR21, plant density ranged from 24 to 35 and 13 to 19 plants m^-2^, respectively. In all experiments, commercial hybrids had the highest plant density; while the Tx430 genotype, an inbred line, had the lowest plant density (Figure S6).

Flowering time was evaluated each year and determined when 50% of the two central rows were flowering at 50%. In AB19 and AB20, one plant from each experimental unit was harvested around flowering to estimate leaf area. In 2019, the leaf area per plant was calculated using LI-3100C (LI-COR, Inc, USA). In AB20, the leaf area and size of each leaf across the canopy profile were estimated using photos processed with the ImageJ program (Rueden et al., 2017). In AB20, photographs of a plant on each experimental unit were taken before flowering. These photographs were printed to measure the leaf angle with a protractor. The angle of the adaxial leaf relative to the stalk was measured in the middle of the canopy. In AB19, AB21, and GR21, the total biomass of a single plant was harvested at physiological maturity to estimate grain weight and harvest index (HI). Similarly, in AB19 and GR21, plant height, panicle length, and panicle exertion were measured on three plants per experimental unit.

### Statistical analysis

Differences between putative traits (non-LT vs. LT) and among sorghum genotypes are expected to occur under high VPD. For this reason, we analyzed records of g_s_ and T_Cirt_ between 12:00 to 16:00 hours (periods with high solar irradiation). Information on T_Cirt_ was recorded from June 23 to August 10 in AB19 and August 11, 15, 18, and 19 in AB20. Each year the variance was analyzed with a mixed model in two stages: first, to test the size effect of the putative trait (eq. 1), and second the size effect of each genotype (eq. 2). The study conducted an additional analysis for genotypes evaluated over all experiments (eq. 3). The models were specified as follows:

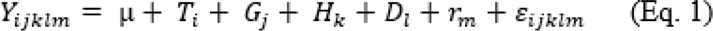

where *Y_ijklm_* is the response variable in the *m^th^* block in the *l^th^*day in the *k^th^* hour in the *j^th^* genotype with the *i^th^* trait, *μ* is the grand mean, *T_i_* is the fixed main effect of the *i^th^* trait, *G_j_* is the random main effect of the *j^th^* genotype, *H_k_* is the random effect of *k^th^* hour, and *r_m_* is the random effect of the *m^th^* replicate.

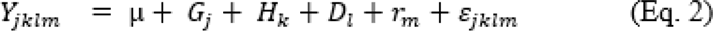

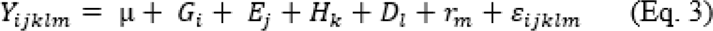

where *Y_jklm_* is the response variable in the *m^th^* replicate in the *l^th^* day in the *k^th^*hour in the *j^th^* genotype, *Y_ijklm_* is the response variable in the *m^th^* block in the *l^th^*day in the *k^th^* hour in the *j^th^* environment with the *i^th^* genotype, *G_i_* is the fixed main of the *i^th^* genotype, *E_j_* is the fixed effect of the *j^th^* environment, *H_k_* is the random effect of *k^th^* hour, and *r_m_* is the random effect of the *m^th^* replicate. A pairwise comparison (Sidak test) was performed when the *F* value fell below α = 0.05 significance threshold.

Broad sense heritability (*H^2^*) was estimated for g_s_, T_Cirt_, and T_Cimg_. *H^2^* was estimated with the lmer library (Bates et al., 2005) by fitting a random effects model which estimated variance components for each random factor on each experiment (Eq. 4) and across experiments (Eq. 5):

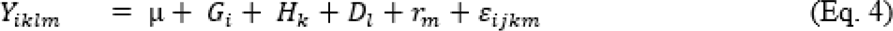

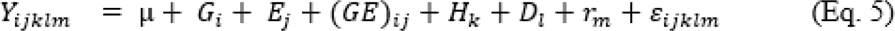

where *Y_iklm_* is the response variable in the *m^th^* replicate in the *l^th^* day in the *k^th^*hour with the *i^th^* genotype, *Y_ijklm_* is the response variable in the *m^th^* replicate in the *l^th^*day in the *k^th^* hour in the *j^th^* environment with the *i^th^* genotype, *G_i_* is the random effect of the *i^th^* genotype, *E_j_* is the random effect of the *j_th_* environment, *(GE)_ij_* is the random two way interaction of *i^th^* genotype *j^th^*environment, *H_k_* is the random effect of *k^th^*hour, and *r_m_* is the random effect of the *m^th^* replicate. The *H^2^* for each experiment and across experiments was calculated with Eq 6 and 7, respectively. The variance for environmental effects *H* and *D* were disregarded in the analysis since they do not contribute to selecting the target phenotype.

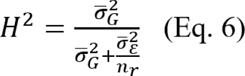

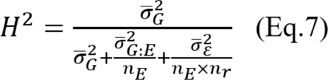

where: *σ*^2^*_G_*, *σ*^2^*_G:E_*, and *σ*^2^_ϵ_ represent variance components of *G*, *G*×*E*, and error, respectively. While,*n*_*E*_ is the number of environments, and *n*_*r*_ is the number of replicates.

Information on g_s_, T_Cirt_, and canopy–air difference was paired with corresponding VPD. Next, g_s_ was analyzed via a linear (Eq. 8), segmented (Eq. 9 and Eq.10), and asymptotic (non-linear, Eq.11) regression analysis.

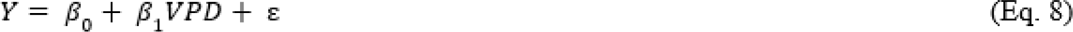

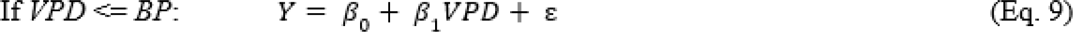

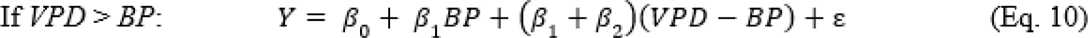

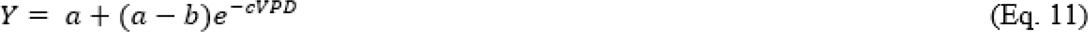

where *Y* represents g_s_, *BP* is the VPD breakpoint, and *β*_0_ is the intercept, *β*_1_ is the slope, and *β*_2_is the second slope, *a* is the plateau (maximum attainable value) for g_s_, *b* is the initial g_s_ when VPD is zero, *c* is the inflexion point or the relative rate of increase for VPD when g_s_ increases. The reason for testing different regressions was to identify the best model that fitted the observed data. Linear regressions were performed using lm library (Bates et al., 2015), the breakpoint regression was analyzed with the library segmented (Muggeo, 2016), non-linear regressions were analyzed with libraries nlme, drc, and aomisc (Ritz et al., 2015). The dependence of g_s_ on VPD was tested via 1000 permutations for linear and breakpoint regressions. To determine this dependence for each genotype, the distribution of the regression coefficient for these permutations was plotted and contrasted against the observed coefficients, *β*_1_for linear regression; *β*_1_, *β*_2_, and BP for segmented regressions. The likelihood-ratio test (library lmtest), and residual standard error (rse) indicated the accuracy of the best regression model (Archontoulis and Miguez, 2015; Kuznetsova et al., 2017).

The effect of canopy architecture traits (i.e., leaf area and leaf size) on transpiration VPD response were visualized using a principal component analysis (PCA) and a correlation analysis. The PCA was estimated with the R prcomp function (“R Core Team,” 2017) and conducted only for AB20 since most canopy architecture traits were evaluated in this experiment. In this analysis, the leaf area represents the maximum leaf area per plant, which occurs around flowering time. The leaf size corresponds to the largest leaf across the canopy profile (Figure S3). The leaf angle is at the attachment between the adaxial leaf and stalk in the central part of the canopy. Plant density represents the number of plants scored after seedling emergence (Figure S6). A correlation analysis was conducted for AB19 and GR21. Grain weight and harvest index for experiments in AB19, AB21, and GR21 were analyzed to characterize agronomic traits of potential donors of putative LT trait.

## RESULTS

### Classification of genotypes for their TR-VPD

To test if VPD during the experiment’s growing seasons represented a typical VPD range, we compared the average daily maximum VPD over 22 years (2000-2021) against the daily maximum VPD recorded on each experiment (Figure 2B and 2C). Results indicate that growing seasons AB19 and AB20 underrepresented average VPD in Kansas; while the growing season in GR21 represented a regular season. In AB19 and AB20, the VPD ranged between 0.2 to 1.9 kPa during the evaluation period (canopy closure to pre-flowering time, Figures 2B). Otherwise in GR21 the VPD ranged between 2.5 to 4.2 kPa.

To determine the existence of genetic variability for TR-VPD in field conditions, we tested i) the putative non-LT versus LT classification and ii) the genotype classification in each experiment (Table 1). To confirm this classification, significance was expected for the fixed effect and differences among mean groups in post hoc analysis. Specifically, higher g_s_ for the putative non-LT trait and higher T_Cirt_ and T_Cimg_ for the LT trait (Figure 1D). Results in all experiments indicated no significance for the trait effect for g_s_, T_Cirt_, and T_Cimg_ (Figure 4, upper panels). However, as expected, the putative non-LT group exhibited slightly high g_s_ (0.06 mol m^-2^ s^-1^) and lower T_Cirt_ (−0.1 C), as indicated in Table S3.

**Figure 4.**
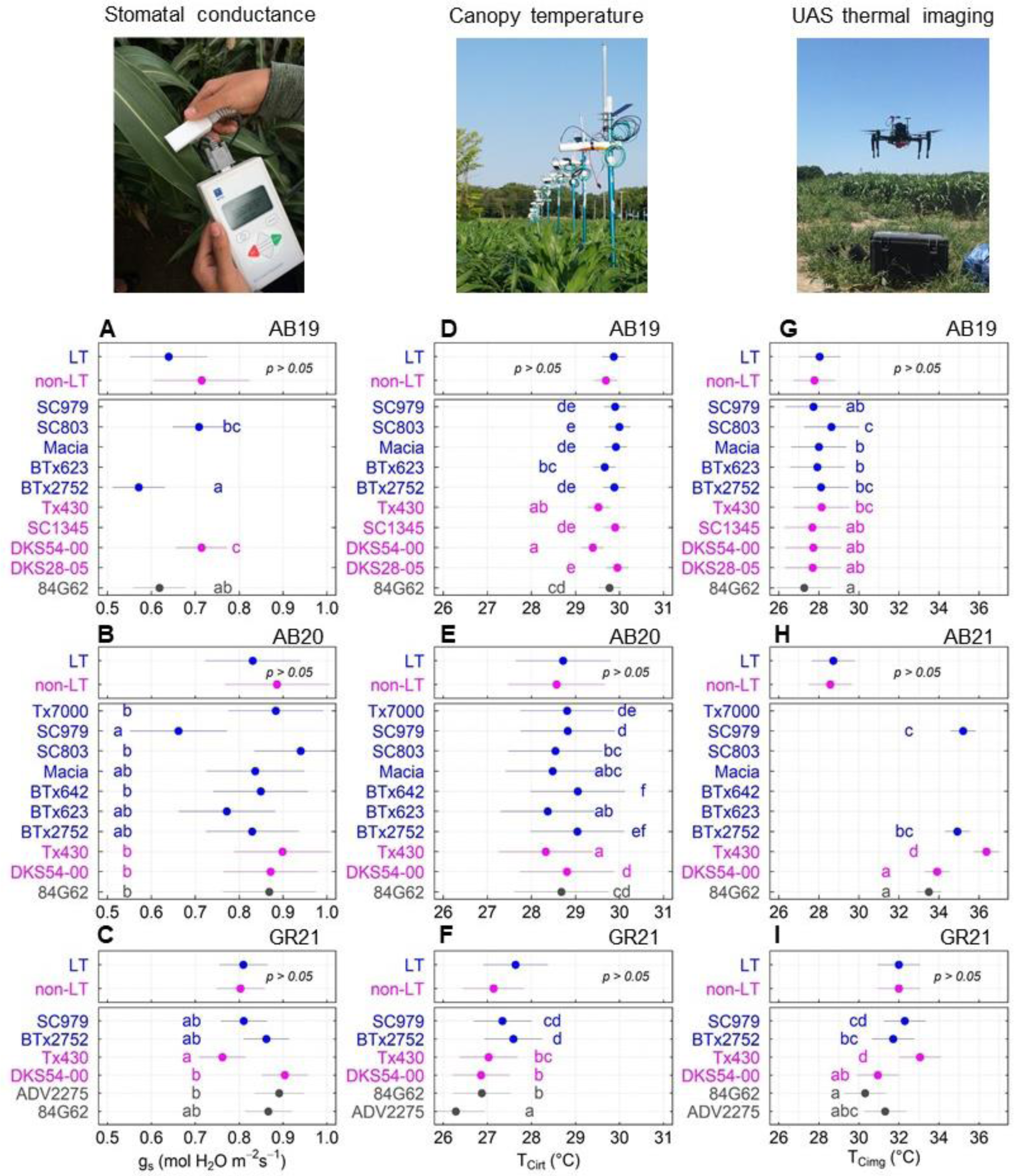
Field Phenomic approaches to discriminate variability in transpiration response to VPD. A, B, and C) Variability for stomatal conductance (g_s_) D, E, and F) Canopy temperature from IRT sensors (T_Cirt_) in hours with high VPD (12:00 to 16:00 hours). H, I and J) Canopy temperature from thermal imagery (T_cimg_) Colors represent the reported putative TR-VPD (Table 1): not known (NN, gray), non-limited transpiration (non-LT, blue) and limited transpiration (LT, magenta). Error lines indicate the standard error. Letters indicate significant differences (α < 0.05) of all pairwise comparisons using the Sidak test. g_s_ represents individual leaves in the sunlit canopy, T_Cirt_ represents the 0.5 m^2^ diameter of the sunlit canopy, while T_Cimg_ represents the median T_C_ at plot level (3 m^2^)

The analysis of variance for each experiment where genotype was the fixed effect indicated differences (*p* < 0.001) for g_s_, T_Cirt_, and T_Cimg_ (Figure 4). In AB19 (Figure 4A, lower panel), as expected, genotypes with putative LT (BTx2752, 0.62 mol m^-2^ s^-1^) and non-LT (DKS54-00, 0.77 mol m^-2^ s^-1^) differed significantly. In this experiment, a genotype classified as LT (SC803) trait unexpectedly exhibited higher g_s_. In AB20 (Figure 4B, lower panel), the mean comparison indicated three groups that differed in g_s_. The first group comprised a genotype with the LT trait SC979, which had the lowest g_s_ (0.73 mol m^-2^ s^-1^). The second group included genotypes with LT (Macia, BTx2752, and BTx2363). The third group had genotypes with non-LT (DKS54-00, Tx430) and LT (Tx7000, BTx642, and SC803) traits, which had the highest g_s_. In GR21 (Figure 4C, lower panel), the genotype Tx430 with the non-LT trait unexpectedly exhibited the lowest g_s_ (Tx430, 0.72 mol m^-2^ s^-1^). As expected, the genotype DKS54-00 with the non-LT trait had the highest (0.90 mol m^-2^ s^-1^) g_s_. Despite genotypes with the LT trait, BTx2752, and SC979, presenting lower g_s_ than DKS54-00, they were not significantly different from this genotype.

In AB19 (Figure 4D, lower panel), phenotypes identified via T_Cirt_ confirmed that six of nine genotypes matched the expectation (Table 1). Genotypes with the LT trait SC979 (29.9 °C), BTx2752 (29.9 °C), Macia (29.9 °C), and SC803 (30.0 °C) had higher T_Cirt_ and differed from genotypes with the non-LT trait DKS54-00 (29.5 °C) and Tx430 (29.6 °C). In AB20 (Figure 4E, lower panel) T_Cirt_ confirmed the classification for Tx7000 (LT), SC979 (LT), BTx642 (LT), BTx2752 (LT), and Tx430 (non-LT). But, it contradicted the classification for genotypes SC803 (LT), Macia (LT), BTx623 (LT), and DKS54-00 (non-LT) (Figure 1D). Genotypes with the LT (BTx2752, 29.0 °C) and non-LT (Tx430, 28.1 °C) traits exhibited the highest and lowest T_Cirt_, respectively. In GR21, all genotypes matched the expectation. The lowest and highest canopy temperature corresponded to genotypes ADV2275 and BTx2752, respectively. In all experiments, T_Cirt_ was replicable only for genotypes with the putative LT trait BTx2752 and SC979.

As expected, in AB21, T_Cimg_ for genotypes with the non-putative-LT trait (DKS54-00, DKS28-05, and SC1345) was lower overall than for genotypes with the putative LT trait (SC803, Macia, BTx623, BTx2752). Discrepant results were obtained for genotype SC979, for which high T_Cimg_ was expected but had low T_Cimg_, and for genotype Tx430 for which low T_Cimg_ was expected but exhibited high T_Cimg_. In AB21 (Figure 4H, lower panel) and GR21 (Figure 4I, lower panel), genotypes SC979 and BTx2752, classified as genotypes with the LT trait, exhibited significantly higher TCimg than genotype DKS54-00, classified as a genotype with the non-LT trait. The only genotype that contradicted its non-LT classification in both experiments was Tx430, which had the highest TCimg.

Over three years of field experiments g_s_ and T_Cirt_ significantly differed (*p* < 0.001), and the covariate effect of VPD on g_s_ was non-significant (Table S5). The pairwise comparison for g_s_ revealed that genotypes DKS54-00 (0.81 mol m^-2^ s^-1^), BTx2752 (0.78 mol m^-2^ s^-1^), and SC979 (0.72 mol m^-2^ s^-1^) belonged to different groups (Table S5). Results of g_s_ for Tx430 (non-LT) were statistically similar to genotypes BTx2752 (LT) and SC979 (LT) (Table S5). The pairwise comparison for T_Cirt_ showed two groups. Genotype DKS54-00 with the non-LT trait had lower T_Cirt_ (28.2 °C) than genotype BTx2752 with the LT trait (28.6 °C). The analysis of T_Cirt_ for four genotypes over three years matched with the TR-VPD classification (Table S5).

To test that different phenomic approaches capture the biology of TR-VPD, we estimated the broad sense of heritability (*H^2^*) and expected similar *H^2^* for g_s_, T_Cirt_, and T_Cimg_. However, H^2^differed among phenomic approaches in each experiment (Figure 5). In AB19, the highest and lowest *H^2^*corresponded for g_s_ (0.5) and T_Cimg_ (0.06). Estimates of *H^2^* Across all experiments for each method indicate a greater *H^2^* for g_s_ (0.4), followed by T_Cimg_ (0.3) and T_Cirt_ (0.2).

**Figure 5.**
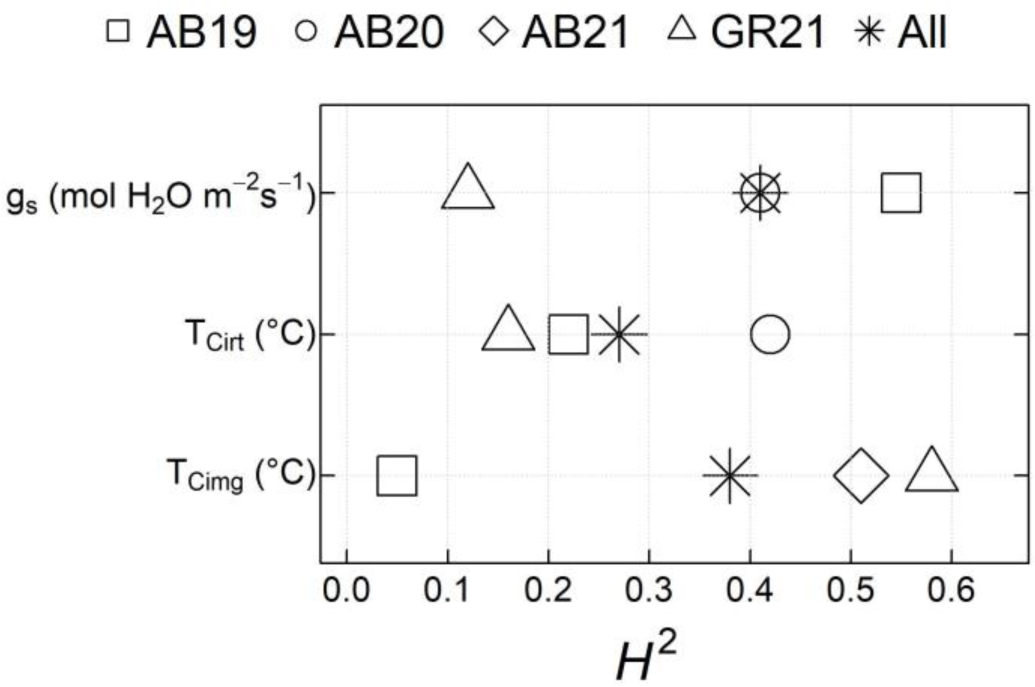
The broad sense of heritability (*H_2_*) for different phenomic approaches to discriminate variability in transpiration response to VPD. Stomatal conductance (g_s_), canopy temperature from IRT sensors (T_Cirt_), and canopy temperature from thermal imagery (T_Cimg_).

### Genotypic variation for stomatal response to VPD

To test that transpiration is driven by VPD and not due to soil water deficit, we compared stomatal response in rainfed and irrigated treatments. This comparison was conducted on a day when the VPD ranged from 3.2 to 4.0 kPa in GR21 (Figure 6A). In this comparison, we expected a significant difference (*p* < 0.05) between treatments, and observations confirmed this expectation. On average, g_s_ under rainfed treatments was 50% lower than in irrigated treatments. In this VPD range, the regression slope for g_s_ in the irrigated treatment was close to zero (0.006 mol m^-2^ s^-1^ kPa); in the rainfed treatment, g_s_ declined (−0.21 mol m^-2^ s^-1^ kPa) as VPD increased. Next, we conducted a regression analysis for each genotype. The stomatal response to VPD was analyzed via linear, segmented, and non-linear models. In all experiments, g_s_ ranged between 0.3 to 1.2 mol m^-2^ s^-1^ and VPD between 1 to 4.5 kPa (Figure 6). The mean g_s_ conductance of 0.79, 0.86, 0.82, and 0.82 mol m^-2^ s^-1^ corresponded to experiments AB19, AB20, AB21, and GR21, respectively. Unexpectedly, high VPD did not increase the mean g_s_ for genotypes with the putative non-LT trait (DKS54-00 and Tx430).

**Figure 6.**
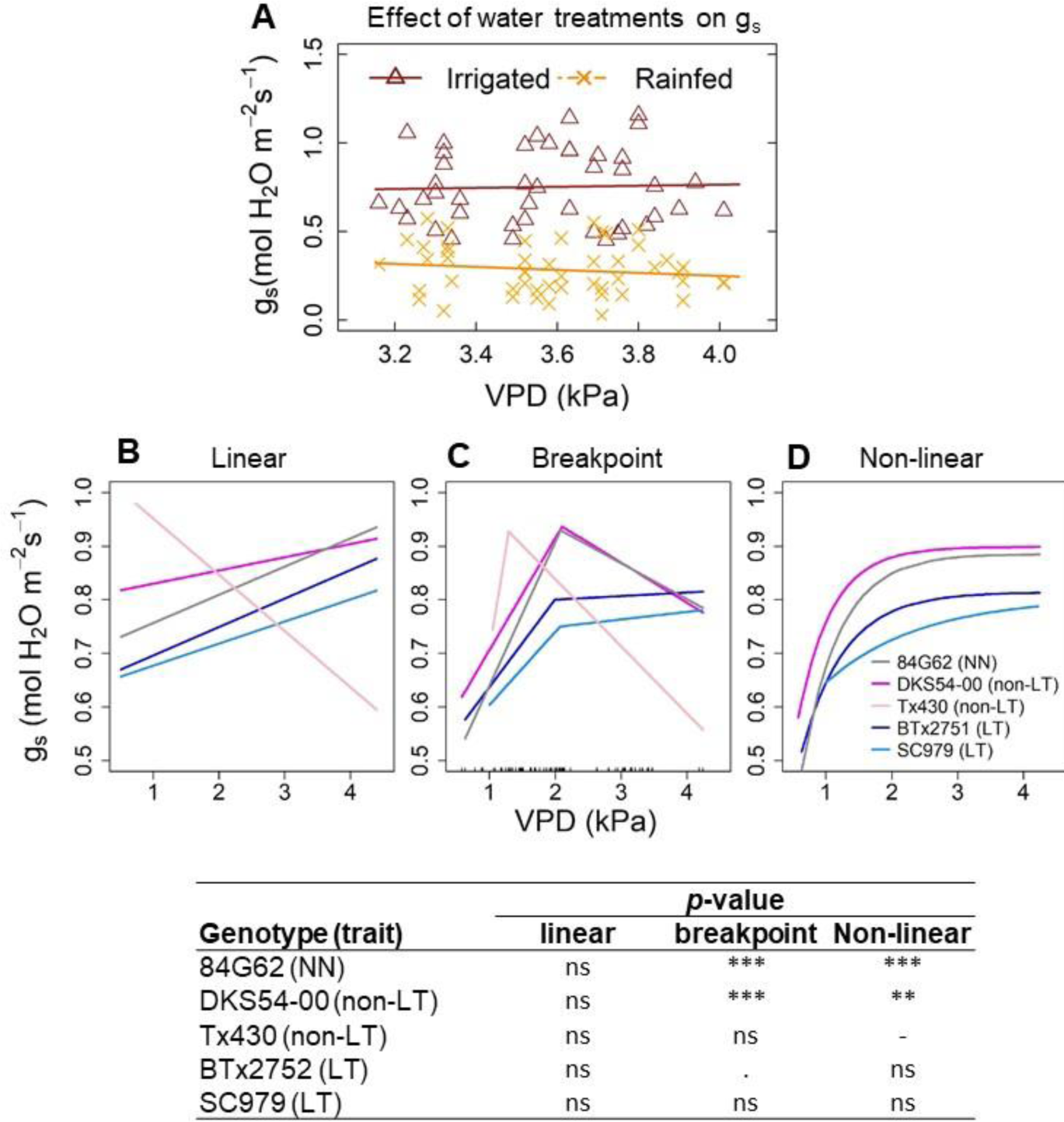
Stomatal response to VPD for sorghum germplasm evaluated over three years of field experiment. A) Comparison of stomatal conductance (g_s_) to VPD under irrigated and rainfed treatments in GR21. The information comprises six genotypes evaluated for their stomatal response to VPD in GR21. B, C and D) variability of stomatal response to VPD for sorghum germplasm represented via linear, breakpoint and non-linear regression models. Each line was fitted with observation indicated in Figure S2.

Linear regression for genotypes 84G62 (NN), DKS54-00 (non-LT), BTx2752 (LT), and SC979 (LT) indicated a positive slope but an unexpected negative one for genotype Tx430 (non-LT) (Figure 6B, Table 3). Segmented (breakpoint) regression for all genotypes suggested a restriction in g_s_ between 1.3 to 2 kPa (Figure 6C). Genotype Tx430, which has a putative non-LT trait, had the lowest breakpoint and the most negative second slope. Breakpoint patterns were similar for commercial hybrids 84G62 (NN) and DKS54-00 (putative non-LT) indicating a negative value for the second slope. Genotypes with the putative LT trait SC979 and BTx2752 exhibited similar patterns, revealing positive values for the first and second slope (Figure 6C). Non-linear regressions revealed an asymptotic curve fitting the observed patterns for all genotypes 84G62, DKS54-00, BTx2752, and SC979, except Tx430. A non-linear regression revealed that differences on transpiration under high VPD are represented via the maximum plateau and the inflection point. Parameters for each regression and genotypes are indicated on Table 3.

To test that the g_s_ is dependent on VPD in linear regression, we compared the observed versus permuted slopes (Figure S3). For genotypes with the non-LT trait DKS54-00 and Tx430, we expected the observed slope to be positive and significantly higher than permuted slopes. For genotypes with the LT trait BTx2752 and SC979, which restrict gas exchange at high VPD, we expected the observed slope to be non-significantly different from the permuted slopes. However, observations contradicted our expectation, indicating that g_s_ and VPD are independent for DKS54-00 (*β*_1_ = 0, *p* = 0.1), while g_s_ depends on VPD, positive slope, for genotypes BTx2752 and SC979 (*β*_1_ = 0, *p* < 0.05). Similarly, a permutation analysis for a breakpoint regression revealed that the regressed variable (g_s_) is independent of the regressor (VPD, Figure S4). To determine the goodness of fit for different regression models, we used the likelihood ratio test. In this analysis, we expected significance for linear regression for genotypes with the non-LT, whereas significance for a breakpoint regression for genotypes with the LT trait. Unexpectedly, breakpoint and non-linear regressions fitted the observed data (*p* < 0.001) for a genotype with non-LT trait DKS54-00 (Figure 6). Otherwise, for a genotype with non-LT trait Tx430, the linear and breakpoint regression revealed no significant difference (*p* > 0.05). Similarly, all regressions were not significantly different for genotypes with LT trait BTx2752 and SC979 (*p* > 0.05, Figure 6).

### Canopy temperature (T_C_) as surrogate method to discriminate TR-VPD

To test if T_C_ is a surrogate method of g_s_, and not an artifact of canopy architecture trais, a PCA was conducted for AB20 and a correlation analysis for AB19 and GR21. In AB20, PC1, PC2 and PC3 represented 38.9, 23.9 and 22.1% of the total variability in the data (Figure 7). PCA1 had a moderately positive loading for leaf area (0.4) and leaf size (0.4) and a negative loading for plant density (−0.4) and T_Cirt_ (−0.4). PCA2 had a positive and negative loading with leaf size (0.5) and g_s_ (−0.6), respectively. PCA3 is dominated by negative loading of the number of leaves (−0.7) and positive loading for T_Cirt_ (0.4). The biplot for PC1 versus PC2 and PC2 versus PC3 revealed that T_Cirt_ and g_s_ are in distinct quadrants suggesting a moderate negative correlation (Figure 7SA) for these phenotypes. Similarly, in GR21, the correlation between g_s_ versus T_Cimg_ was negative and significant (*r* = −0.9, *p* < 0.05). A biplot PC2 versus PC3 showed a substantial correlation between T_Cirt_ and the number of leaves. These results indicate that T_Cirt_ is a good proxy of g_s_, but canopy architecture can affect T_C_. Otherwise, biplots suggested a strong correlation among canopy architecture traits. For instance, leaf area negatively correlated with the number of leaves (PC1 versus PC2) and leaf angle (PC2 versus PC3).

**Figure 7.**
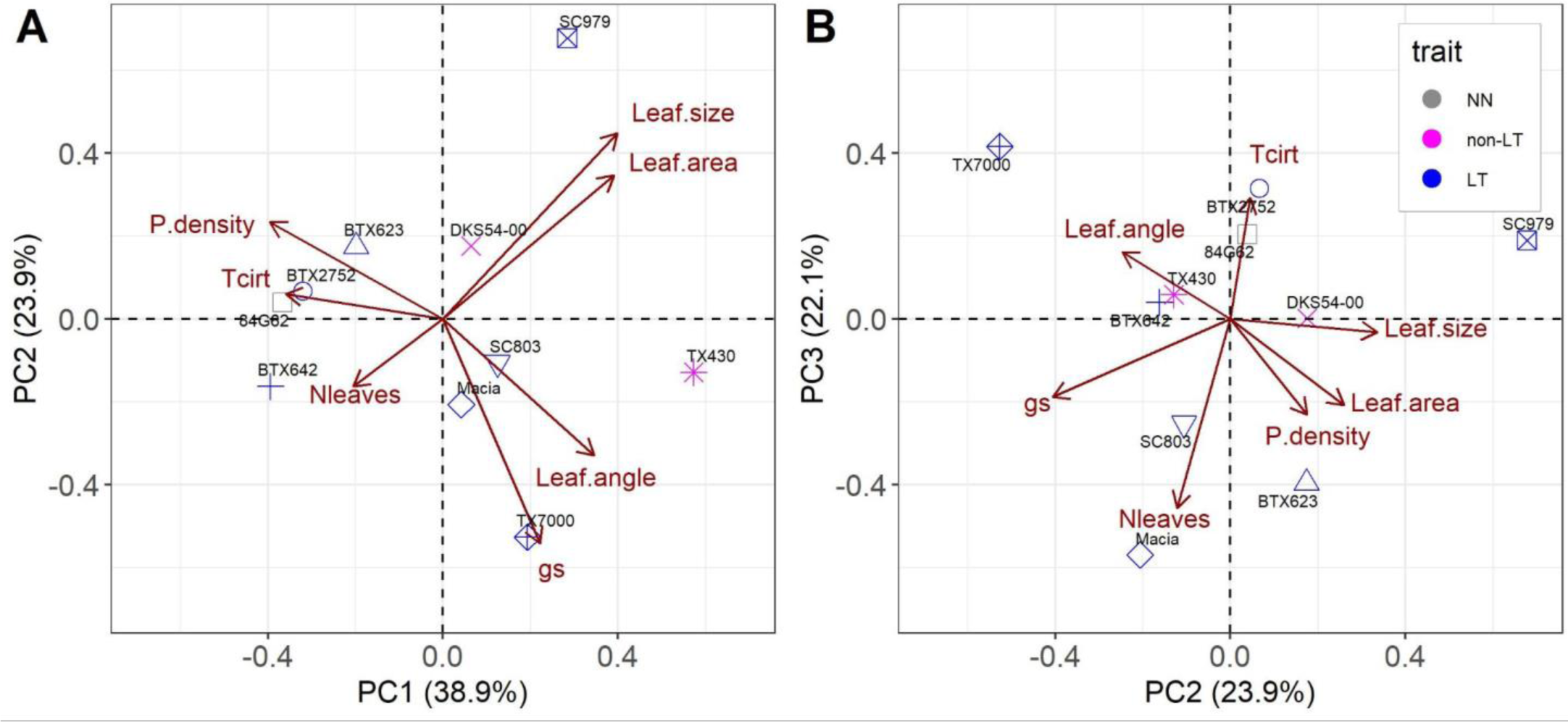
Principal component analysis (PCA) for genotypes characterized for their TR-VPD and canopy architecture traits in AB20. P.density: plant density (plants m^-2^); Leaf.angle: leaf angle (°); Leaf.area: leaf area per plant (leaf area plant^-1^); Leaf.size: maximum leaf size (cm^2^); N.leaf: maximum number of leaves. Details on each canopy architecture trait is provided in Table S4 and Figures S3, S5, and S6.

To test if TR-VPD phenotype is an artifact of stand count, we conducted correlations between g_s_, T_Cirt_, T_Cimg_, and plant density (Figure S7). In AB20, the correlation was close to zero for plant density and T_Cirt_, while a negative correlation (−0.5) between plant density and g_s_ was non-significant (*p* > 0.05). Significant negative correlations between plant density and T_Cimg_ (Figure S7B and S7C) were obtained in AB19 (*r* = −0.8, *p* < 0.01) and GR21 (*r* = 0.9, *p* < 0.05). Likewise, in GR21 (Figure S7C), the correlation between plant density and g_s_ was significant (*r* = 0.9, *p* < 0.01). Overall, results suggest that stand count is the main trait that can hamper the effectiveness of any phenomic approach when dissecting TR-VPD.

## DISCUSSION

Testing the existence of genetic variability for the putative LT trait via different phenomic approaches is pivotal when deciding to include this trait in a breeding pipeline. This is the first study that phenotyped the TR-VPD in field trials in sorghum production regions of the United States. Our findings i) help to better understand the challenges when testing the replicability of the trait via different phenomic approaches, ii) discuss the best representation of genetic diversity by comparing different regression models, iii) review the underlying physiological mechanism of the putative LT trait and iv) outline future steps for crop breeding when using the LT to deliver water efficient sorghum hybrids.

### A phenotype can be similar or vary among phenomic approaches

Our study confirmed the TR-VPD classification via three phenomic approaches for three out of eleven genotypes (Figure 4). Results for genotypes SC979 (LT), BTx2752 (LT), and DKS54-00 (non-LT) corresponded with prior references (Gholipoor et al., 2010; Shekoofa et al., 2014). A PCA (Figure 7) and a negative correlation between g_s_ and T_C_, although non-significant, to some extent, suggested T_C_ as a surrogate phenotype to discriminate differences in transpiration (Figure 7, Figure S7). The effectiveness of T_C_ in determining variability for LT transpiration has been demonstrated by comparing T_C_ with whole-plant transpiration in controlled environments (Belko et al., 2013).

For other genotypes, the g_s_ phenotype contradicted prior classification (Figure 4). For instance, unexpected high g_s_ for genotypes with the putative LT trait SC803, Tx7000, and BTx642 (Table 1) suggest they had the non-LT trait. Discrepant results were also reported for lentils where a genotype with the putative LT in the greenhouse displayed a non-LT trait in field trials (Guiguitant et al., 2017). These observations imply that growth chamber experiments do not fully represent field conditions. Phenotyping for TR-VPD in growth chambers is limited to early growth stages and environmental conditions are more erratic in field settings. Indeed, prior studies suggested that early growth stages in field conditions (Shekoofa et al., 2014) and temperatures beyond 30 °C (Riar et al., 2015) suppress the expression of the LT trait.

The phenotype identified with each phenomic approach varied for most genotypes (Figure 4). These differences can be attributed to the sample size utilized in each method (Figure 4), variability in canopy architecture, and stand count. The interaction of canopy architechture traits with wind speed influences convective heat transfer and T_C_ (Gates, 1980; Leigh et al., 2017; Melcher et al., 1994). Canopy architecture traits can be more important determinants of T_C_ than physiological traits (Still et al., 2021; Woods et al., 2018) since canopy architechture traits and wind speed influence convective heat transfer and T_C_. Genotypes with an acute leaf angle likely disrupted air movements through the canopy, leading to high T_Cirt_, as observed in genotype BTx642 (Figure 7A). Soil exposure in plots of genotypes with fewer leaves would have reflected solar energy, contributing to increasing air temperature and T_C_ (Figure 7B). This would explain the unexpectedly high T_Cirt_ in AB19 for genotypes DKS28-00 and SC1345 with the putative non-LT trait and low leaf area (Table S6).

Microenvironmental conditions created by low plant density and high tillering can override the phenotypic expression of a target trait (Jones, 2007). The phenotype identified via g_s_ for genotype Tx430 shifted from non-LT (Figure 4A, 4B) to LT (Figure 4C) when the plant density was lower than 13 pl m^-2^ (Figure S8A). Although the experiment was well-irrigated, at low plant density, it is possible that soil evaporation surpassed plant transpiration. High tillering (Figure S8B) likely exacerbated water demand, promoting water competition among culms (Borrell et al., 2014), leading to low g_s_ and high T_Cimg_ (Figures 4H and 4I).

T_Cimg_, an HTPP approach, can accelerate breeding selection by bringing higher precision (Deery et al., 2019), if such an approach can capture the nature of the target trait. Nevertheless, *H^2^*for T_Cimg_ was close to zero and lower than *H^2^* for g_s_ in AB19. While the *H^2^*for g_s_ (0.16-0.55) in our study aligned with an investigation for cotton (0.16 to 0.44) (Percy et al., 1996), the *H^2^* for T_Cimg_ ranged from 0.05 to 05, which are lower than the values reported for wheat (0.3 to 0.8) (Anderegg et al., 2021; Deery et al., 2019). Different *H^2^* for T_Cimg_ (Figure 5) among experiments indicates that this approach is prone to microenvironmental conditions likely caused by the variability of canopy architecture traits (Figure 7 and S7A). Additionally, stand count can override the expression of the TR-VPD (Figure S7B and S7B). Overall, using T_Cimg_ to screen large breeding populations would require considering the number of leaves and stand count as covariate effects.

### Non-linear function models better represent genetic variability for TR-VPD

Genetic variability identified via different phenomic approaches in this study provides candidate donor lines to breed sorghum yields for water deficit conditions (Figure 4). Our study has shown that genetic variability occurs under low and high (<2 kPa) VPD (Figure 6D). Genetic variability for TR-VPD has been proposed as differences in slopes and restriction (breakpoint) in gas exchange (Gholipoor et al., 2010; Shekoofa et al., 2014). In our study, observations indicated that no genotypes with the non-LT trait increased g_s_ (Figure S3). Evidence of whether observed data fit a linear or segmented regression was never shown or discussed in prior studies. For instance, observed data for outdoor pots suggested asymptotic patterns for chickpeas with LT and non-LT traits, but the study reported a breakpoint and LT and linear response (Zaman-Allah et al., 2011).

In our research, stomatal response to VPD fit different regressions (Figure 6), indicating the breakpoint or asymptotic as the best model for a genotype with non-LT trait. Although disparities between a breakpoint (Figure 6C) and an asymptotic model (Figure 6D) seem to be negligible, independent data support that at higher VPD, g_s_ will reach its maximum value and remain constant (Figure 6A). Our findings align with studies where transpiration under well-watered conditions and high VPD remained constant. A wheat study reported breakpoints response for 100% of commercial varieties tested in growth chambers (Schoppach et al., 2017). For woody and herbaceous plants, transpiration under controlled environments reached a maximum plateau at a VPD of 2.5 kPa (Turner et al., 1984). An asymptotic function better portrays the stomatal response to VPD, as this non-linear model has a biological meaning and is utilized to describe photosynthesis, light intensity, and light interception (Archontoulis and Miguez, 2015). The correct mathematical representation of TR-VPD has a significant implication when modeling the effect of the LT at crop system scales. Simulation studies using segmented and nonlinear functions are most likely overestimating the impact of the LT trait on harvested yield.

### The underlying mechanism of TR-VPD

The underlying physiological mechanism of TR-VPD remains enigmatic. The LT trait has been associated with aquaporin inhibition (Maurel et al., 2016) and low hydraulic conductivity (Choudhary et al., 2013b) that causes partial stomatal closure. In arabidopsis and angiosperms, stomatal response to VPD is controlled by hydropassive and hydroactive processes (Merilo et al., 2018; Wang et al., 2001). Nevertheless, studies in soybean and peanut claim that LT is modulated via a hydropassive mechanism (Sinclair et al., 2017). While the hydropassive response involves changes in the turgor cell, the hydroactive mechanism is modulated via metabolic signaling (Merilo et al., 2018; Wang et al., 2001). Studies reported leaf-derived ABA when plants are exposed to high VPD (Hu et al., 2016; Jalakas et al., 2021), although findings indicate that *Open Stomata 1* (*OST1*) kinase rather than ABA controls stomatal response to humidity (Merilo et al., 2018). Otherwise, differences in leaf hydraulic conductivity reported for genotypes SC1205 (dwarf, non-LT) and SC15 (tall, putative LT) (Choudhary et al., 2013b; Ocheltree et al., 2013) can result from contrasting hydraulic anatomy (Du et al., 2020) and plant architecture. Indeed, under well-water conditions, a study reported differences in hydraulic anatomy (xylem diameter) for a short (ICSSH58, 1.5 m) and tall (ICSV25280, 1.7 m) sorghum (Guha et al., 2018).

Discrepancies in TR-VPD between non-LT and LT traits are hypothetically due to stomatal openness (Sinclair et al., 2017) with expected open stomata for the genotype with the non-LT trait and partial stomatal closure for a genotype with the LT trait (Figure 8A). Determining whether partial stomatal closure is controlled by hydropassive or hydroactive mechanisms requires testing the expression of metabolic signals (Figure 8B). Otherwise, asymptotic patterns of TR-VPD (Figure 6D) suggest that the stomata remain open at high VPD for genotypes with LT and non-LT traits (Figure 8C). Then, differences in gas exchange are hypothetically due to variations in hydraulic anatomy traits such as variability in stomatal density (Bheemanahalli et al., 2021), xylem diameter, or boundary layer thickness (Figure 8D). From a breeding perspective, a hydroactive mechanism would allow identifying known genes such as *OST1* or *slow anion channel 1* (*SLAC1*), which are ABA-dependent or -independent (Merilo et al., 2018). Meanwhile, the hydropassive mechanism implies that the aquaporin response is independent of any signaling pathway and unknown genes. Testing these hypotheses in identified lines (DKS54-00, BTX2752, and SC979) can elucidate mechanisms underlying the TR-VPD. Further quantifying the carbon assimilation trade-offs of the LT trait in near-isogenic lines would indicate the impact of this trait on drought-prone regions. This knowledge can guide breeding programs to consider different options when designing a breeding pipeline.

**Figure 8.**
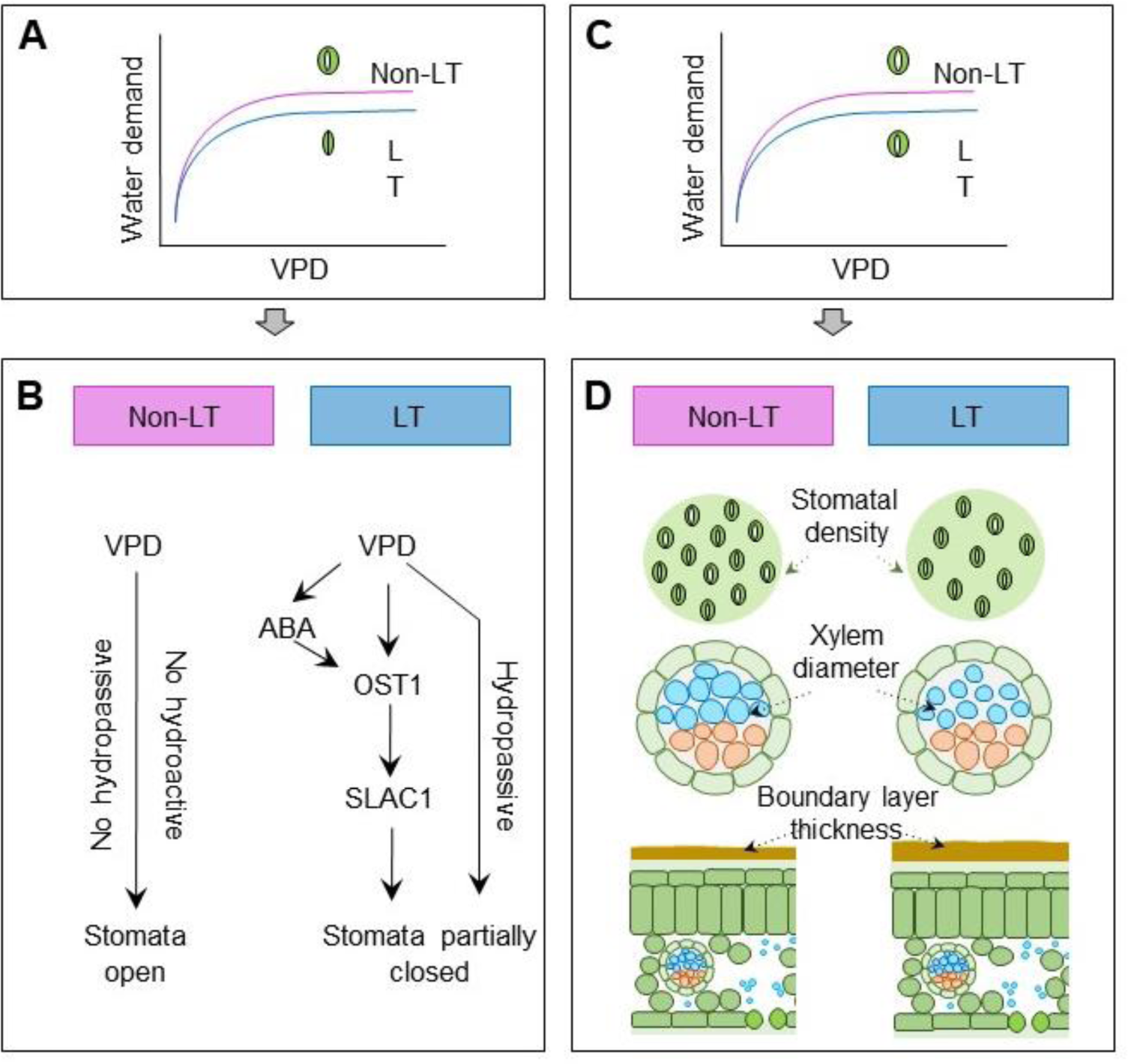
Hypotheses on mechanisms underlying variation in transpiration VPD response. A) Water demand varies due to stomatal response, where the genotype with putative non-LT maintains open stomata, and the genotype with LT trait partially closes the stomata. B) Both hydroactive and hydropassive mechanisms can cause stomatal closure, but hormonal signaling is involved in the hydroactive mechanism. C) Genotypes with non-LT (pink line) and LT (blue) maintain stomata open, and water demand varies due to differences in hydraulic anatomy. D) Expected hydraulic anatomic traits for genotypes with putative non-LT and LT traits. The hydropassive and hydroactive mechanism for the LT trait was adapted from Merilo et al. (2018).

### Future steps

The results of this study represent TR-VPD when soil water is not limited in the soil profile. However, drought-prone regions are subjected to simultaneous high VPD and soil water deficit. Results under a rainfed treatment and soil water deficit in a semiarid environment revealed a decline in g_s_ as VPD increases (Figure 6A), suggesting that soil water deficit and high VPD lead to stomatal closure to avoid dehydration and physiological damage (Oren et al., 1999). So far, the confounding effect of VPD and soil moisture has been ignored in studies of LT. Then it would be worthwhile to determine whether the effect of VPD on stomatal closure prevails over the impact of soil water deficit on stomatal closure or vice versa. Genotypes with putative LT traits identified in this study correspond to full-season backgrounds. A comparison of yield components under well-watered and rainfed treatments in a semiarid environment (GR21) demonstrated that early flowering time overrode the effect of the putative LT trait, either in irrigated or rainfed treatments (Figure S10). In silico simulations showed that the LT trait can increase yields by more than 5% in western regions of the sorghum belt where medium and short-season hybrids are planted. Then it would be needed to identify and introgress the LT trait in short and medium-season backgrounds. Developing a mapping population and near-isogenic lines with donors SC979, BTx2752 would allow identifying QTLs via T_Cimg_; nevertheless, confounding effects detailed in this study need to be considered when using this approach.

**Table 2.** Regression parameters for linear, breakpoint and non-linear stomatal conductance (g_s_) response to VPD for five genotypes. Information represents data collected in AB19, AB20, AB21, and GR21. 84G62:176 observations, DKS54-00: 173 observations, Tx430: 133 observations, Btx2752: 175 observations, and Sc979: 128 observations. RSE: residual standard error.

| Genotype | intercept | slope |  |  | R<sup>2</sup> | RSE |
| --- | --- | --- | --- | --- | --- | --- |
| *Linear* | $\text{mol m}^{-2} \text{s}^{-1}$ | $\text{mol m}^{-2} \text{s}^{-1} \text{kPa}^{-1}$ | | | | |
| 84G62 (NN) | $0.70 \pm 0.04$ | $0.05 \pm 0.01$ | | | 0.03 | 0.23 |
| DKS54-00 (non-LT) | $0.80 \pm 0.04$ | $0.02 \pm 0.02$ | | | 0.00 | 0.27 |
| Tx430 (non-LT) | $0.98 \pm 0.06$ | $-0.08 \pm 0.02$ | | | 0.00 | 0.24 |
| BTx2752 (LT) | $0.64 \pm 0.04$ | $0.05 \pm 0.02$ | | | 0.04 | 0.24 |
| SC979 (LT) | $0.80 \pm 0.04$ | $0.02 \pm 0.02$ | | | 0.00 | 0.25 |
| *Breakpoint* | intercept<br>$\text{mol m}^{-2} \text{s}^{-1}$ | slope1<br>$\text{mol m}^{-2} \text{s}^{-1} \text{kPa}^{-1}$ | slope2<br>$\text{mol m}^{-2} \text{s}^{-1} \text{kPa}^{-1}$ | BP<br>kPa | R<sup>2</sup> | RSE |
| 84G62 (NN) | $0.37 \pm 0.07$ | $0.26 \pm 0.04$ | $-0.06 \pm 0.04$ | $2.08 \pm 0.18$ | 0.16 | 0.21 |
| DKS54-00 (non-LT) | $0.49 \pm 0.08$ | $0.20 \pm 0.05$ | $-0.07 \pm 0.03$ | $2.10 \pm 0.20$ | 0.09 | 0.22 |
| TX430 (non-LT) | $-0.057 \pm 1.42$ | $0.76 \pm 1.21$ | $-0.12 \pm 0.29$ | $1.29 \pm 0.20$ | 0.11 | 0.24 |
| BTx2752 (LT) | $0.47 \pm 0.04$ | $0.16 \pm 0.09$ | $0.00 \pm 0.02$ | $1.99 \pm 0.45$ | 0.06 | 0.21 |
| SC979 (LT) | $0.46 \pm 0.16$ | $0.13 \pm 0.09$ | $0.01 \pm 0.04$ | $2.07 \pm 0.56$ | 0.01 | 0.21 |
| *Asymptotic* | plateau<br>$\text{mol m}^{-2} \text{s}^{-1}$ | inflexion point<br>$\text{mol m}^{-2} \text{s}^{-1} \text{kPa}^{-1}$ | start | | | RSE |
| 84G62 (NN) | $0.89 \pm 0.02$ | $1.47 \pm 0.40$ | $-0.00 \pm 0.36$ | | | 0.22 |
| DKS54-00 (non-LT) | $0.89 \pm 0.02$ | $1.96 \pm 0.68$ | $-0.09 \pm 0.60$ | | | 0.23 |
| Tx430 (non-LT) | - | - | - |  |  | - |
| BTx2752 (LT) | $0.81 \pm 0.03$ | $1.52 \pm 0.81$ | $0.03 \pm 0.62$ | | | 0.21 |
| SC979 (LT) | $0.48 \pm 0.06$ | $0.69 \pm 1.86$ | $0.48 \pm 0.62$ | | | 0.24 |

## ACKNOWLEDGMENT

This study was supported by funding from the Foundation for Food and Agriculture Research - Seeding Solution “CA18-SS-0000000094 – Bridging the Genome-to-Phenome Breeding Gap for Water-Efficient Crop Yields (G2P Bridge)” to G.P.M. and S.S.B.; the Kansas Department of Agriculture “Collaborative Sorghum Investment Program Water Optimized Sorghum for Kansas” to G.P.M; and the Kansas Grain Sorghum Commission.

## AUTHOR CONTRIBUTIONS

G.P.M., S.S.B., T.F., and R.R. contributed to the conception and design of the work. T.F. and R.R. collected field experimental data. X.W was in charge of the UAS imaging. R.R. and X.W. conducted data analysis, interpretation, and drafting of the article. G.P.M., S.S.B., T.F., R.R., X.W., J.P., A.E.L. contributed to the final manuscript.

