## Supplemental Tables for "Field phenomics reveals genetic variation for transpiration response to vapor pressure deficit in sorghum"

### SUPPLEMENTARY INFORMATION

**Table S1. Field management and environmental conditions for experiments conducted in Ashland Bottoms (AB) Kansas from 2019 (19) to 2021(21) and in Greeley (GR) Colorado in 2021 (21).**

| Experiment name | AB19 | AB20 | AB21 | GR21 |
|---|---|---|---|---|
| Latitude | 39.14309 | 39.13891 | 39.13891 | 40.44724 |
| Longitude | -96.63194 | -96.64055 | -96.64055 | -104.63695 |
| Soil texture | Clay | Clay | Clay | Sandy loam |
| Initial soil nutrient | 30-20-50 | 60-50-50 | - | 80-50-20 |
| N fertilization | 170 kg $ha^{-1}$ | 180 kg $ha^{-1}$ | - | 200 kg $ha^{-1}$ |
| Experimental unit | 4 row-plot | 8 row-plot | 8 row-plot | 8 row-plot |
| Seeds per row | 60 | 80 | 80 | 80 |
| Water supply | Drip lines | Drip lines | | Subsur. Irrigat |
| Planting date | 3-Jun | 13-Jun | 11-Jun | 09-Jun |
| Emergence date | 11-Jun | 20-Jun | 25-Jun | 19-Jun |
| Flowering date | 10-Aug | 20-Aug | 24-Aug** | 19-Aug*** |
| Maturity date | 05-Oct | - | 04-Oct | - |
| Harvest date | 24-Oct | 11-Nov | - | 22-Nov |
| CGDD* | 1444 | 1448 | 1560 | 1145 |
| Tmax (C) | 29.8 | 29.2 | 31.4 | 30 |
| Tmin (C) | 18.7 | 16.4 | 18 | 11 |
| Rain (mm) | 520 | 413 | 338 | 107 |
| VPD (kPa) | 1.2 | 1.4 | - | 3.1 |
| Soil moisture ($m^3 m^{-3}$) | 0.3 | 0.25 | | 0.2 |

- Not available
* Cumulative growing degree days.
** Flowering time did not vary in well watered and rainfed treatment.
*** Indicates flowering time for irrigated treatments. Flowering time for the rainfed treatment occurred four days later. Although not all genotypes bloomed in this environment.
Note: Information on flowering time for each genotype is indicated in Table S7.

**Table S2. Environmental characteristics during evaluations of stomatal conductance (gs) and thermal imageries in field experiments in Kansas and Colorado.**

| **Experiment** | **Date** | **Hours** | **AirVPD (kPa)** | **Solar rad. (W m$^{-2}$)** | **Tmax (°C)** | **Soil moisture* (m$^3$ m$^{-3}$)** |
|---|---|---|---|---|---|---|
| *Stomatal conductance (gs)* | | | | | | |
| AB19 | 03-Aug-19 | 13 - 16 | 1.1 | 1039 | 28.8 | 0.31 - 0.12 |
| AB19 | 04-Aug-19 | 9 - 13 | 1.2 | 950 | 28.8 | 0.31 - 0.12 |
| AB19 | 06-Aug-19 | 10 - 14 | 1.7 | 927 | 31.1 | 0.28 - 0.12 |
| AB19[r] | 13-Aug-19 | 10 -12 | 1.3 | 825 | 30.2 | 0.31 - 0.26 |
| AB19[r] | 19-Aug89 | 10 - 16 | 1.6 | 865 | 33.6 | 0.30 - 0.27 |
| AB19[r] | 29-Aug-19 | 13 - 16 | 1.8 | 808 | 32.9 | 0.28 - 0.27 |
| AB20 | 01-Aug-20 | 12 - 14 | 1.5 | 987 | 28.0 | 0.28 - NA |
| AB20 | 09-Aug-20 | 13 - 15 | 1.3 | 687 | 29.8 | 0.22 - NA |
| AB20 | 14-Aug-20 | 12 - 14 | 1.6 | 973 | 32.5 | 0.20 - NA |
| AB21 | 09-Aug-21 | 14 - 16 | 2.1 | 804 | 34.3 | 0.20 -0.21 |
| AB21 | 11-Aug-21 | 14 - 16 | 2.0 | 775 | 34.5 | 0.20 - 0.21 |
| AB21 | 12-Aug-21 | 14 - 15 | 2.2 | 896 | 34.9 | 0.21 - 0.19 |
| AB21 | 18-Aug-21 | 14 - 16 | 1.9 | 879 | 31.6 | 0.20 - 0.19 |
| AB21 | 25-Aug-21 | 14 - 16 | 2.1 | 776 | 33.8 | 0.24 - 0.20 |
| AB21 | 26-Aug-21 | 14 - 16 | 3.2 | 765 | 36.3 | 0.24 - 0.20 |
| AB21 | 27-Aug-21 | 14 - 15 | 2.6 | 777 | 34.2 | 0.24 - 0.20 |
| GR21 | 28-July-21 | 13 - 15 | 4.2 | 924 | 35.0 | 0.26 - NA |
| GR21 | 29-July-21 | 14 - 16 | 4.2 | 836 | 34.8 | 0.26 - NA |
| GR21 | 10-Aug-21 | 14 - 15 | 3.5 | 848 | 31.6 | 0.24 - NA |
| GR21 | 11-Aug-21 | 13 - 15 | 3.3 | 886 | 31.2 | 0.26 - NA |
| GR21 | 18-Aug-21 | 12 - 14 | 3.1 | 887 | 32.0 | 0.26 - NA |
| GR21 | 27-Aug-21 | 13 - 15 | 3.6 | 881 | 32.3 | 0.27 - NA |
| *UAV-thermal imagery data collection* | | | | | | |
| AB19 | 26-Jul-19 | 10 | 1.1 | 1039 | 28.8 | 0.31 - 0.12 |
| AB19 | 26-Jul-19 | 12 | 1.1 | 1039 | 28.8 | 0.31 - 0.12 |
| AB19 | 26-Jul-19 | 14 | | | | |
| AB21 | 06-Aug-21 | 14 | | | | |
| AB21 | 09-Aug-21 | 14 | 2.1 | 804 | 34.3 | 0.20 -0.21 |
| GR21 | 17-Aug-21 | 14 | 3.1 | 887 | 32.0 | 0.26 - NA |

*Soil moisture at 30 cm and 15 cm.

[r] Evaluations during reproductive stage, not included in the analysis

Table S3. Components of the linear mixed model for hypothetical classification of non-LT and LT traits for canopy IRT and stomatal conductance in AB19 and AB20.

| | Canopy IRT - 2019 | | | Canopy IRT - 2020 | | |
|---|---|---|---|---|---|---|
| *Predictors* | *Estimates* | *CI* | *p* | *Estimates* | ***CI*** | ***p*** |
| (Intercept) | 29.8 | 29.2 – 30.4 | <0.001 | 28.7 | 26.8 – 30.6 | <0.001 |
| trait [non-LT] | -0.1 | -0.4 – 0.1 | 0.183 | -0.1 | -0.6 – 0.3 | 0.541 |
| *Random Effects* | | | | | | |
| $\sigma^2$ | 0.46 | | | 0.26 | | |
| $\tau_{00\ genotype}$ | 0.04 | | | 0.08 | | |
| $\tau_{00\ hour}$ | 0.09 | | | 0.23 | | |
| $\tau_{00\ date}$ | 0.70 | | | 3.46 | | |
| $\tau_{00\ rep}$ | 0.00 | | | 0.00 | | |
| ICC | 0.64 | | | 0.94 | | |
| $N_{genotype}$ | 9 | | | 8 | | |
| $N_{hour}$ | 5 | | | 5 | | |
| $N_{date}$ | 12 | | | 4 | | |
| $N_{rep}$ | 3 | | | 3 | | |
| Observations | 1415 | | | 428 | | |
| Marginal $R^2$ / Conditional $R^2$ | 0.006 / 0.647 | | | 0.001 / 0.936 | | |

| | Conductance - 2019 | | | Conductance - 2020 | | |
|---|---|---|---|---|---|---|
| *Predictors* | *Estimates* | *CI* | *p* | *Estimates* | *CI* | *p* |
| (Intercept) | 624.2 | 403.7 – 844.6 | <0.001 | 844.8 | 592.2 – 1097.4 | <0.001 |
| trait [non-LT] | 71.2 | -158.4 – 301.0 | 0.543 | 68.7 | -57.8 – 195.1 | 0.287 |
| *Random Effects* | | | | | | |
| $\sigma^2$ | 13747.3 | | | 36201.6 | | |
| $\tau_{00\ genotype}$ | 8656.9 | | | 4487.0 | | |
| $\tau_{00\ hour}$ | 39939.5 | | | 5022.3 | | |
| $\tau_{00\ date}$ | 174.51 | | | 37102.2 | | |
| $\tau_{00\ rep}$ | | | | 6585.0 | | |
| ICC | 0.78 | | | 0.60 | | |
| $N_{genotype}$ | 3 | | | 8 | | |
| $N_{hour}$ | 5 | | | 4 | | |
| $N_{date}$ | 3 | | | 3 | | |
| $N_{rep}$ | | | | 4 | | |
| Observations | 87 | | | 177 | | |
| Marginal $R^2$ / Conditional $R^2$ | 0.017 / 0.784 | | | 0.010 / 0.599 | | |

**Table S4. Components of the linear mixed model for stomatal conductance and canopy IRT in sorghum genotypes in AB19, AB20 and GR21.**

| | Conductance | | | Canopy IRT | | |
|---|---|---|---|---|---|---|
| ***Predictors*** | ***Estimates*** | ***CI*** | ***p*** | ***Estimates*** | ***CI*** | ***p*** |
| (Intercept) | 992.2 | 702.6 – 1281.8 | <0.001 | 27.9 | 26.1 – 29.7 | <0.001 |
| genotype [BTX2752] | -48.1 | -108.1 – 11.8 | 0.116 | 0.6 | 0.4 – 0.9 | <0.001 |
| genotypeDKS54-00 | 49.6 | -10.1 – 109.4 | 0.103 | 0.2 | -0.0 – 0.5 | 0.083 |
| genotype [SC979] | -109.6 | -178.7 – -40.5 | 0.002 | 0.5 | 0.3 – 0.8 | <0.001 |
| genotype [TX430] | -131.4 | -200.3 – -62.6 | <0.001 | 0.2 | -0.04 – 0.5 | 0.101 |
| kPa.VPD | -94.2 | -183.8 – -4.5 | 0.039 | | | |
| Random Effects | | | | | | |
| $\sigma^2$ | 58442.51 | | | 0.70 | | |
| $\tau_{00\ date}$ | 13398.84 | | | 1.29 | | |
| $\tau_{00\ hour}$ | - | | | 0.06 | | |
| $\tau_{00\ rep}$ | 330.04 | | | 0.01 | | |
| $\tau_{00\ year}$ | 35341.72 | | | 2.31 | | |
| ICC | 0.46 | | | 0.84 | | |
| $N_{date}$ | 12 | | | 24 | | |
| $N_{year}$ | 3 | | | 3 | | |
| $N_{hour}$ | - | | | 5 | | |
| $N_{rep}$ | 4 | | | 3 | | |
| Observations | 550 | | | 1122 | | |
| Marginal $R^2$ / Conditional $R^2$ | 0.131 / 0.528 | | | 0.012 / 0.842 | | |

-Not applicable

**Table S5. Stomatal conductance and canopy IRT and for sorghum genotypes.** Each value represents the great mean over three experiments (AB19, AB20 and GR21) in Kansas (KS) and Colorado (CO). Hypothetical TR-VPD for each genotype are not known (NN), non-limited transpiration (non-LT) and limited transpiration (LT). Letters indicate significant differences (α < 0.05) of all pairwise comparisons using the Sidak test.

| **Genotype (trait)** | **Conductance** **mol $m^{-2}$ $s^{-1}$ kPa** | | **Canopy IRT** **C** | **Groups** |
|---|---|---|---|---|
| 84G62 (NN) | 0.81 ± 0.04 | bc | 27.9 ± 0.93 | a |
| DKS54-00 (non-LT) | 0.87 ± 0.04 | c | 28.2 ± 0.93 | a |
| Tx430 (non-LT) | 0.77 ± 0.04 | ab | 28.1 ± 0.93 | a |
| BTx2752 (LT) | 0.78 ± 0.04 | b | 28.6 ± 0.93 | b |
| SC979 (LT) | 0.72 ± 0.04 | a | 28.5 ± 0.93 | b |

| Source of variation | df | *p* | df | *p* |
|---|---|---|---|---|
| Genotype | 4 | <0.001 | 4 | <0.001 |
| VPD | 1 | 0.24 | - | - |

- Not applicable.

Note: When the Tx430 genotype is part of the analysis, the interaction of genotype by VPD is significant. However, removing the genotype from the analysis makes this interaction negligible.

**Table S6. Canopy architecture traits in AB19 and AB20.** Each value represents the mean over four replications. The leaf size, leaf length and leaf width corresponds to the largest leaf of the canopy profile. The canopy profile for each genotype is presented in Figure S3A.

| | AB19 | | AB20 | | | | |
|---|---|---|---|---|---|---|---|
| Genotype | N leaves | Leaf area $m^2$ $plant^{-1}$ | Leaf area $m^2$ $plant^{-1}$ | Leaf size $cm^2$ | Leaf length cm | Leaf width cm | Leaf angle ° |
| SC803 | 17 ± 1 | 4.1 ± 0.2 | 3.7 ± 0.5 | 510 ± 51 | 82.4 ± 3.2 | 8.1 ± 0.8 | 34.8 ± 12.9 |
| SC979 | 15 ± 1 | 4.6 ± 0.5 | 4.5 ± 0.4 | 658 ± 45 | 96.3 ± 2.3 | 8.8 ± 0.4 | 33.3 ± 10.2 |
| BTx2752 | 17 ± 1 | 3.4 ± 0.5 | 2.2 ± 0.3 | 426 ± 65 | 85.1 ± 3.6 | 6.5 ± 0.6 | 33.0 ± 7.3 |
| BTx623 | 15 ± 1 | 3.7 ± 0.6 | 3.2 ± 0.3 | 467 ± 32 | 88.9 ± 0.8 | 7.1 ± 0.2 | 26.4 ± 2.3 |
| BTx642 | - | - | 2.9 ± 0.1 | 334 ± 8 | 69.7 ± 3.0 | 6.5 ± 0.3 | 26.4 ± 8.4 |
| Macia | 16 ± 1 | 3.9 ± 0.3 | 3.7 ± 0.4 | 469 ± 41 | 79.1 ± 1.1 | 7.9 ± 0.4 | 33.8 ± 7.0 |
| TX7000 | - | - | 2.4 ± 0.5 | 415 ± 45 | 77.8 ± 1.8 | 7.6 ± 0.4 | 40.8 ± 10.5 |
| SC1345 | 14 ± 1 | 3.7 ± 0.6 | - | - | - | - | - |
| Tx430 | 16 ± 2 | 4 ± 1.1 | 4 ± 0.7 | 567 ± 76 | 90.7 ± 3.1 | 7.9 ± 0.8 | 36.3 ± 6.9 |
| *DKS28-05* | *11 ± 2* | *2.4 ± 0.1* | - | - | - | - | - |
| *DKS54-004* | *15 ± 1* | *4.0 ± 0.2* | *3.7 ± 0.7* | *508 ± 39* | *86.2 ± 2.8* | *7.6 ± 04* | *29.5 ± 9.5* |
| *84G62* | *16 ± 1* | *3.6 ± 0.9* | *2.8 ± 0.6* | *440 ± 64* | *79.7 ± 0.9* | *7.2 ± 0.7* | *30.0 ± 9.5* |

- Not applicable

**Table S7. Flowering time represented in Julian days and cumulative thermal time for sorghum genotypes in Kansas and Colorado**. Dae: days after emergence. GDD: growing degree days. Each value represents four replications per genotype.

| Genotype | AB19 | | AB20 | | AB21 | | GR21 | |
|---|---|---|---|---|---|---|---|---|
| | Dae day | GDD C | Dae day | GDD C | Dae day | GDD C | Dae day | GDD C |
| *Irrigated* | | | | | | | | |
| SC803 | 60 | 869 | 60 ± 2 | 928 ± 21 | - | - | - | - |
| SC979 | 60 | 869 | 61 ± 1 | 942 ± 7 | 59 ± 7 | 924 ± 128 | 68 ± 2 | 833 ± 18 |
| BTx2752 | 65 ± 6 | 948 ± 91 | 64 ± 2 | 984 ± 27 | 59 ± 6 | 929 ± 99 | 81 ± 2 | 974 ± 21 |
| BTx623 | 60 | 869 | 66 ± 4 | 1014 ± 58 | - | - | - | - |
| BTx642 | - | - | 68 ± 2 | 1056 ± 28 | - | - | - | - |
| Macia | 63 ± 5 | 909 ± 79 | 62 ± 3 | 939 ± 78 | - | - | - | - |
| Tx7000 | - | - | 59 ± 2 | 918 ± 21 | - | - | - | - |
| SC1345 | 52 ± 4 | 752 ± 52 | 53 ± 2 | 841 ± 39 | - | - | - | - |
| Tx430 | 63 ± 5 | 909 ± 79 | 63 ± 3 | 960 ± 52 | 60 ± 5 | 944 ± 95 | 77 ± 3 | 936 ± 24 |
| *DKS28-05* | *48 ± 1* | *676 ± 3* | *50 ± 1* | *779 ± 18* | - | - | - | - |
| *DKS54-004* | *59 ± 3* | *847 ± 45* | *60 ± 2* | *928 ± 21* | *57 ± 7* | *885 ± 122* | *68 ± 1* | *837 ± 15* |
| *84G62* | *59 ± 3* | *847 ± 45* | *59 ± 2* | *921 ± 28* | *59 ± 5* | *915 ± 95* | *75 ± 3* | *916 ± 24* |
| *ADVG2275* | | | | | *58 ± 5* | *914 ± 90* | *67 ± 2* | *821 ± 28* |
| *Rainfed* | | | | | | | | |
| SC979 | - | - | - | - | 60 ± 6 | 943 ± 103 | 74 ± 3 | 899 ± 34 |
| BTx2752 | - | - | - | - | 60 ± 5 | 945 ± 71 | *No fl.* | *No fl* |
| Tx430 | - | - | - | - | 56 ± 1 | 861 ± 83 | 80 ± 1 | 962 ± 12 |
| *DKS54-004* | - | - | - | - | *56 ± 7* | *873 ± 68* | *74 ± 5* | *904 ± 55* |
| *84G62* | - | - | - | - | *58 ± 7* | *912 ± 327* | *No fl* | *No fl* |
| *ADVG2275* | - | - | - | - | *58 ± 5* | *908 ± 84* | *77 ± 4* | *933 ± 34* |

-Not available

No fl: no flowering

**Table S8. Yield components for sorghum genotypes in Kansas (AB) and Colorado (GR)**. The yield represents two central rows per plot. Each value is the mean of four replications. Grain represents the weight of a panicle. HI represents the ratio of grain to biomass. The mean

| Genotype | AB19 | | | AB21 | | GR21 | |
|---|---|---|---|---|---|---|---|
| | **Yield Mg ha$^{-1}$** | **Grain gr plant$^{-1}$** | **HI** | **Grain gr plant$^{-1}$** | **HI** | **Grain gr plant-1** | **HI** |
| *Irrigated* | | | | | | | |
| SC803 | 6.9 ± 0.1 | 41 ± 22 | 0.4 ± 0.2 | - | - | - | - |
| SC979 | 6.7 ± 0.3 | 31 ± 9 | 0.3 ± 0.1 | 53 ± 13 | 0.2 ± 0.1 | 29 ± 7 | 0.4 ± 0.4 |
| BTx2752 | 6.8 ± 0.5 | 31 ± 10 | 0.3 ± 0.2 | 58 ± 16 | 0.3 ± 0.1 | 0 | 0 |
| BTx623 | 6.8 ± 0.1 | 44 ± 11 | 0.5 ± 0.1 | - | - | - | - |
| Macia | 6.6 ± 0.6 | 47 ± 26 | 0.4 ± 0.2 | - | - | - | - |
| SC1345 | 6.4 ± 0.2 | 41 ± 2 | 0.4 ± 0.1 | - | - | - | - |
| Tx430 | 4.9 ± 0.7 | 18 ± 11 | 0.2 ± 0.2 | 47 ± 25 | 0.2 ± 0.1 | 0 | 0 |
| *DKS28-05* | *6.5 ± 0.1* | *46 ± 9* | *0.5 ± 0.1* | - | - | - | - |
| *DKS54-004* | *7.2 ± 0.3* | *37 ± 8* | *0.5 ± 0.1* | *62 ± 18* | *0.3 ± 0.1* | *56 ± 18* | *0.5 ± 0.2* |
| *84G62* | *7.3 ± 0.1* | *62 ± 19* | *0.5 ± 0.1* | *77 ± 12* | *0.3 ± 0.1* | *36 ± 17* | *0.3 ± 0.2* |
| *ADVG2275* | - | - | - | *78 ± 11* | *0.3 ± 0.1* | *60 ± 9* | *0.6 ± 0.1* |
| *Rainfed* | | | | | | | |
| SC979 | - | - | - | 41 ± 14 | 0.2 ± 0.1 | 9 ± 5 | 0.1 ± 0.1 |
| BTX2752 | - | - | - | 46 ± 16 | 0.3 ± 0.1 | 0 | 0 |
| Tx430 | - | - | - | 28 ± 18 | 0.1 ± 0.1 | 0 | 0 |
| *DKS54-004* | - | - | - | *46 ± 27* | *0.2 ± 0.1* | *29 ± 11* | *0.5 ± 0.2* |
| *84G62* | - | - | - | *62 ± 23* | *0.3 ± 0.1* | *2 ± 3* | *0.1 ± 0.1* |
| *ADVG2275* | - | - | - | *50 ± 12* | *0.3 ± 0.1* | *14 ± 13* | *0.2 ± 0.3* |

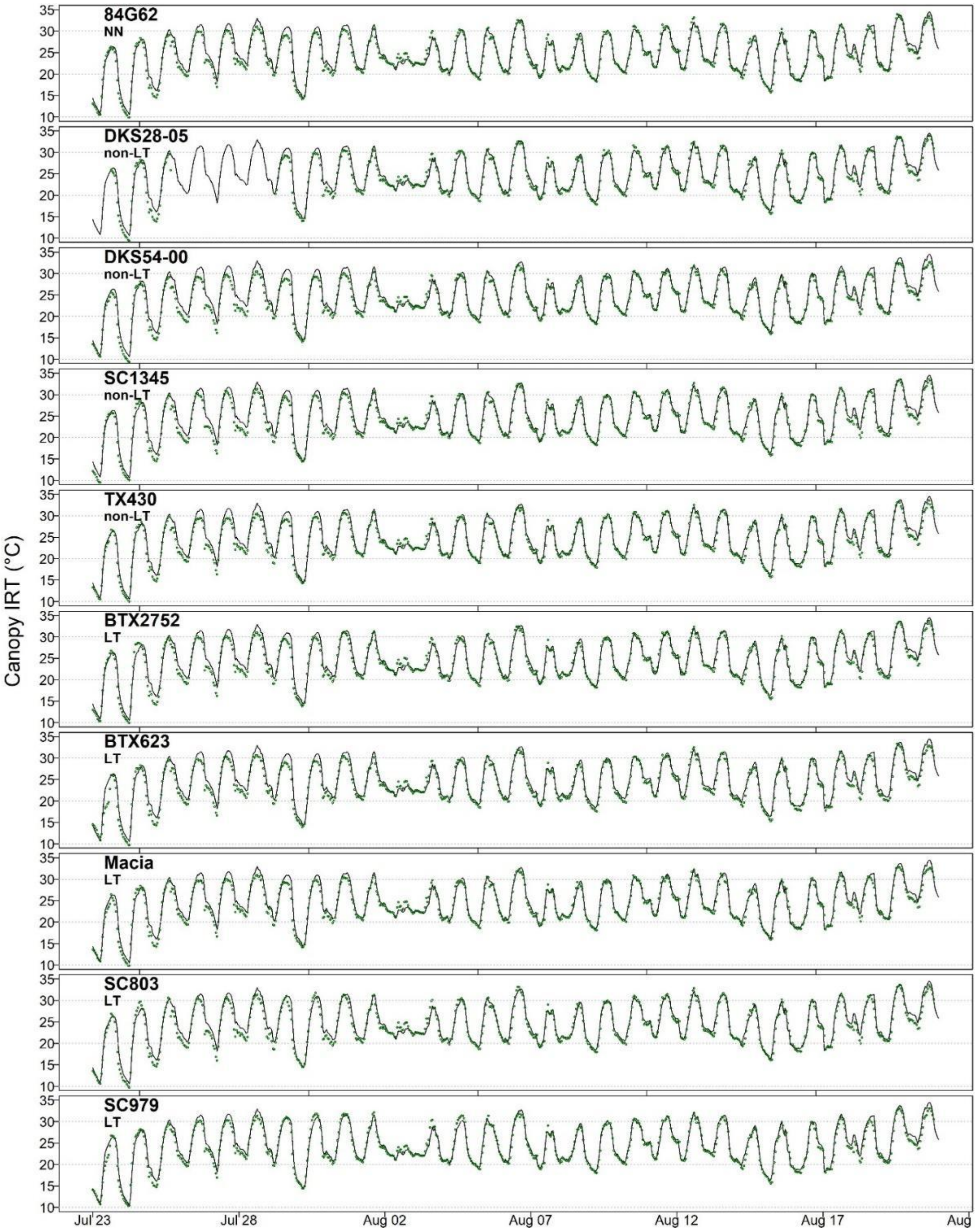


Figure S1. Hourly air temperature (black solid lines) and canopy IRT (green dots) for sorghum genotypes in Ashland bottoms, Kansas 2019 (AB19). Each green point represents the mean of three records. NN: not known, non-LT: non-limited transpiration trait, and LT: limited transpiration trait.

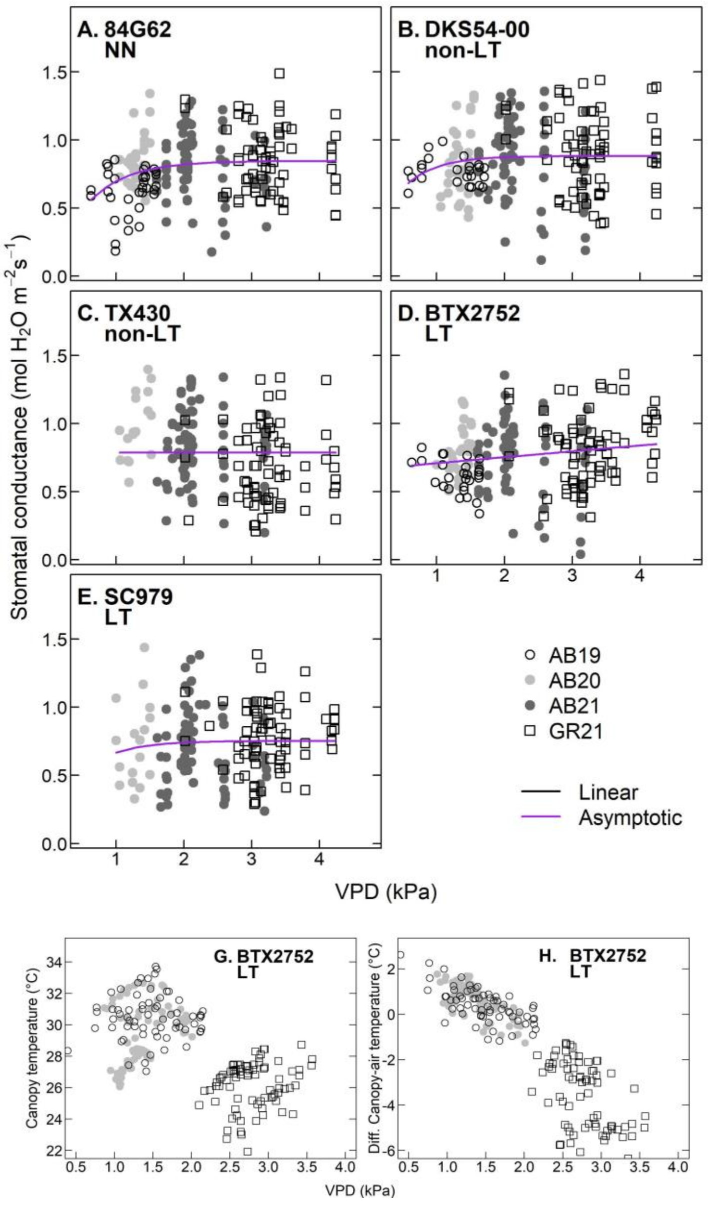


**Figure S2. Stomatal conductance and canopy temperature response to VPD for sorghum germplasm evaluated over three years of field experiment**. A-E) Stomatal conductance response for genotypes with putative non-LT and LT traits under well-watered conditions. The black and purple lines indicate linear and non-linear responses. G) Canopy temperature response to VPD for a genotype with putative LT trait. H) Canopy air-temperature difference for a genotype with putative LT trait.

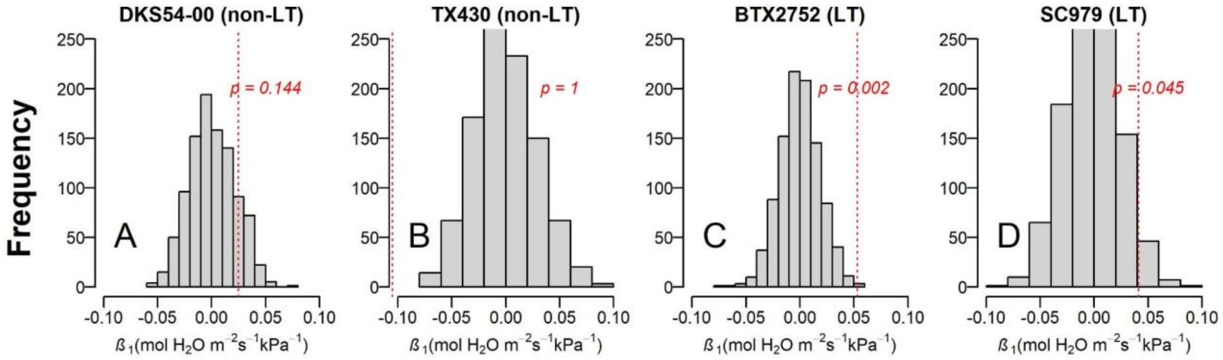


Figure S3. Distribution of the $\beta_1$ for linear regressions (Eq. 9) generated via 1000 permutations for each genotype. Null hypothesis ($H_o$): $\beta_1$ = 0 and alternative hypothesis ($H_a$): $\beta_1 > 0$. $H_o$ is rejected when *p*-value < 0.05, indicating that $g_s$ depends on VPD. The dashed red line indicates $\beta_1$ for the original regression.

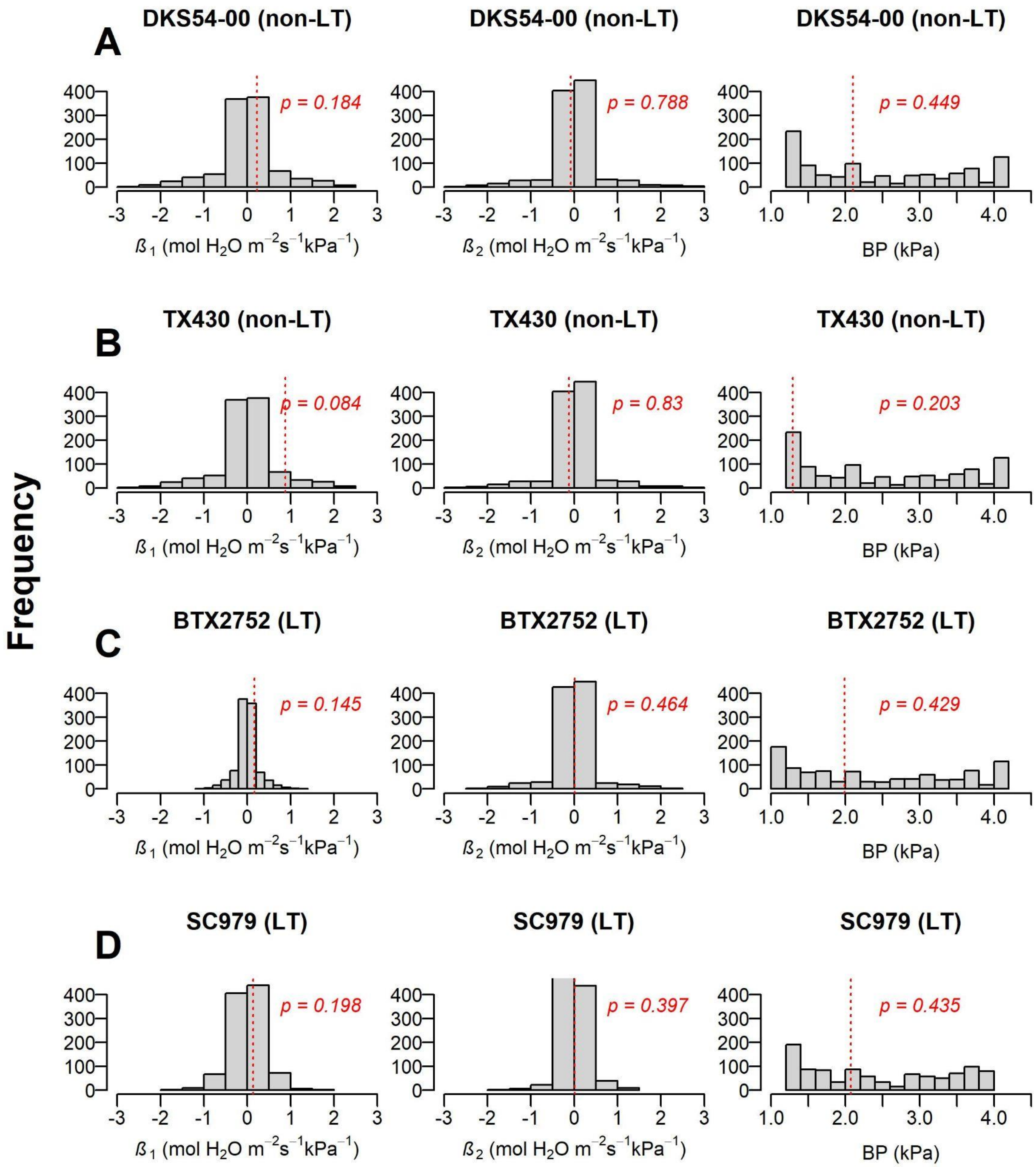


Figure S4. Distribution of the $\beta_1$, $\beta_2$ and BP for segmented regressions (Eq. 9) generated via 1000 permutations for each genotype. Null hypothesis ($H_o$): $\beta_1$ = 0, $\beta_2$ = 0, BP = 0 and alternative hypothesis ($H_a$): $\beta_1$ > 0, $\beta_2$ < 0, BP > 0 . $H_o$ is rejected when *p*-value < 0.05, indicating that $g_s$ depends on VPD. The dashed red line indicates $\beta_1$, $\beta_2$ and BP for the original regression.

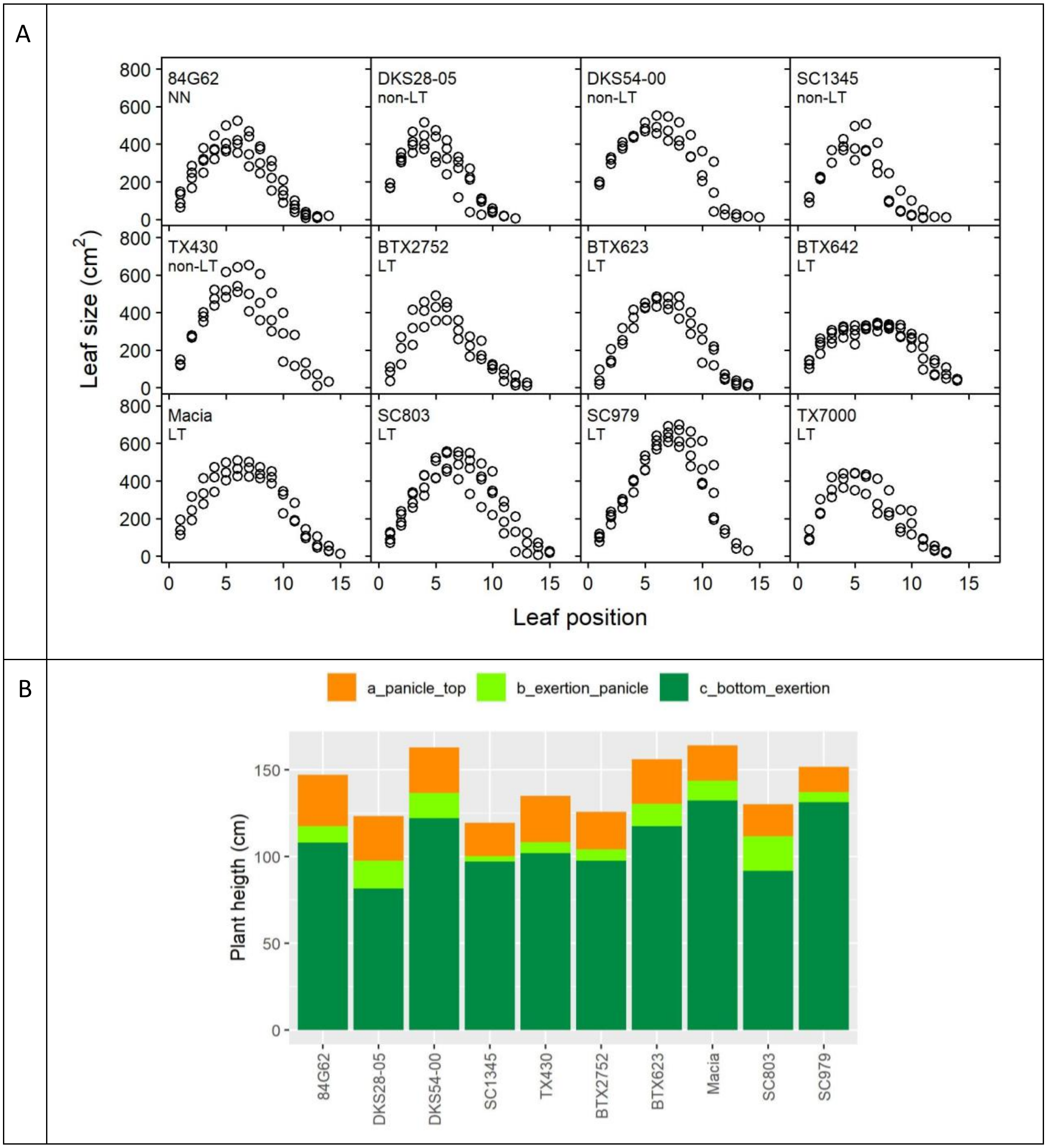


**Figure S5. Canopy architecture of sorghum genotypes characterized for their transpiration response to VPD.** A) Canopy profile for 12 sorghum genotypes in Ashland Bottoms, 2020 (KS). Leaf number one is the flag leaf. NN: not known, non-LT: non limited transpiration trait, and LT: limited transpiration trait. Each panel represents the canopy profile of three plants that were harvested after flowering time. B) Plant height, panicle length and exertion length for 10 genotypes in Ashland Bottoms, 2019 (KS). Each bar plot represents the mean of nine plants that were evaluated at the hard dough stage.

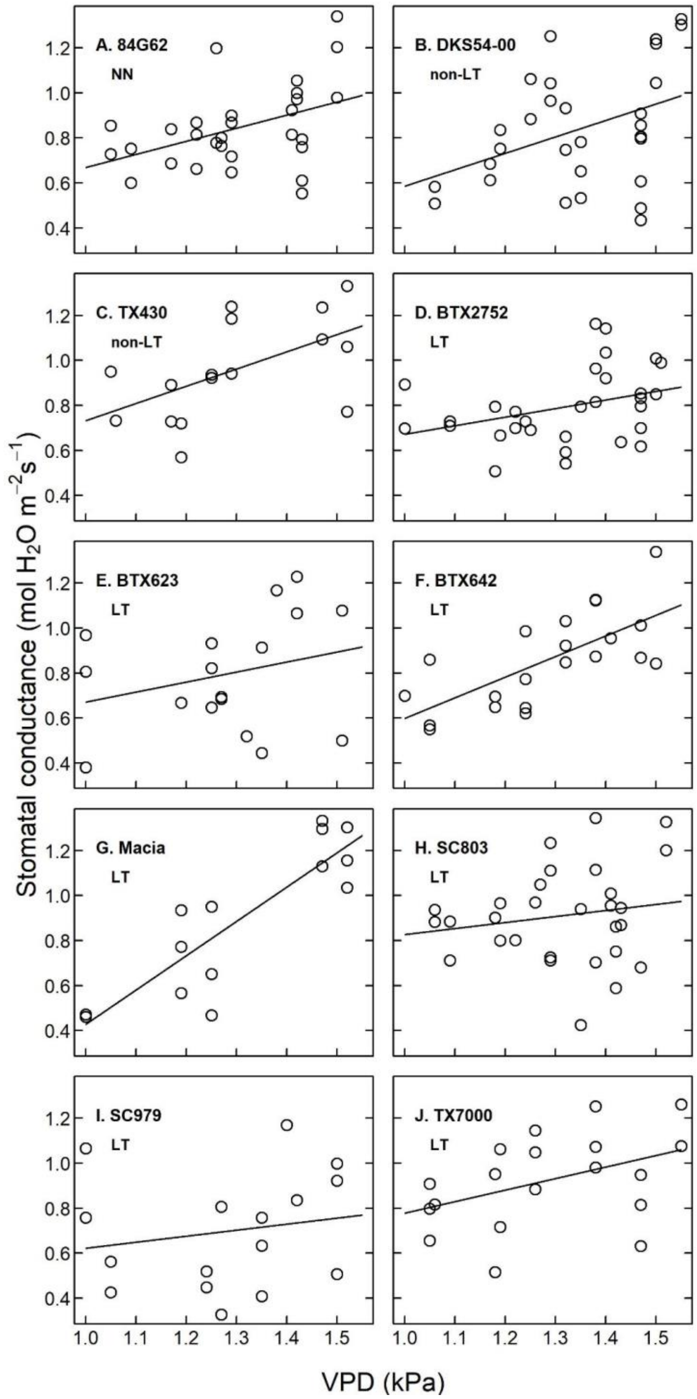


Figure S6. Stomatal conductance response to VPD in sorghum genotypes in AB20 (KS). NN: not known, non-LT: non limited transpiration trait, and LT: limited transpiration trait.

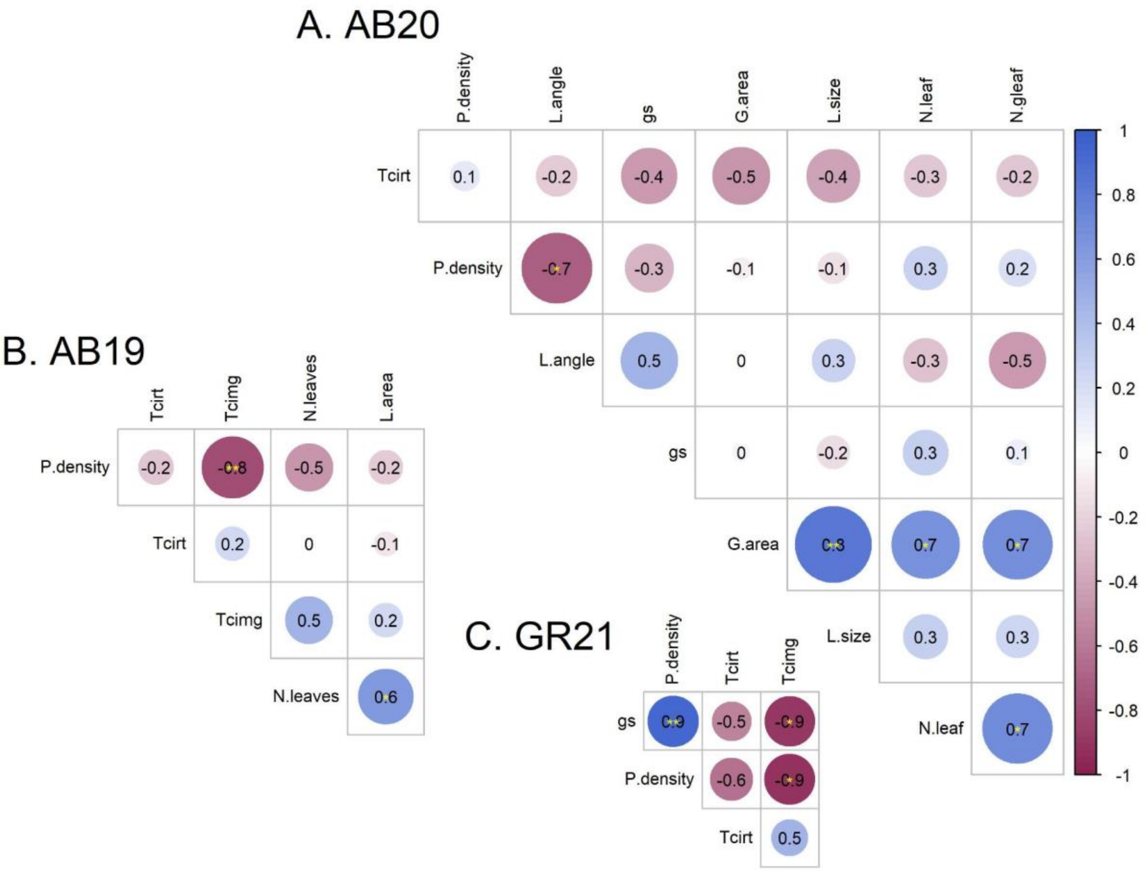


**Figure S7. Correlation of transpiration response to VPD and canopy architecture traits in sorghum germplasm.** A and B) Correlations for experiments in AB20 and AB19, Kansas. C) Correlation for experiments in GR21, Colorado. Values in each circle represent the correlation; asterisk indicates the p-value significance of $\alpha < 0.05$ (*), $\alpha < 0.01$ (**), and $\alpha < 0.001$(***). Density: plant density (plants $m^{-2}$); L.angle: leaf angle (°); G.area: green leaf area per plant (leaf area $plant^{-1}$); L.size : maximum leaf size ($cm^2$); N.leaf: maximum number of leaves. Information on each canopy architecture trait is provided in Table S4, Figure S4, and Figure S6.

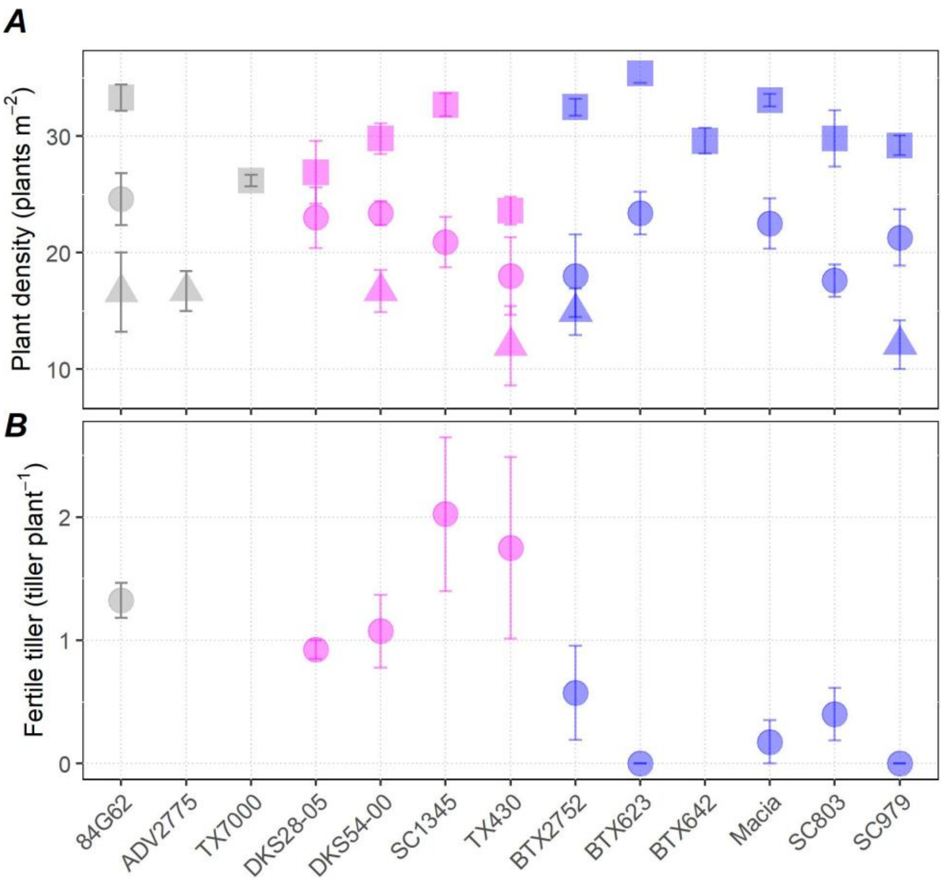


**Figure S8. Plant density and number of fertile tillers for experiments in AB19, AB20, and GR21.** Solid circles represent AB19, solid squares represent AB20, and solid triangles represent GR21. Each point represents the mean of four points and the vertical lines represent the standard error. Colors represent the reported putative TR-VPD (Table 1): not known (NN, gray), non-limited transpiration (non-LT, pink) and limited transpiration (LT, blue).
